# Plant cells tolerate high rates of organellar mistranslation

**DOI:** 10.1101/2025.03.07.642005

**Authors:** Benjamin Brandt, Sebastian Schwartz, Serena Schwenkert, Moritz Krämer, Kuenzang Om, Carina Engstler, Andreas Klingl, Peter Jahns, Jürgen Eirich, Iris Finkemeier, Asaph Benjamin Cousins, Hans-Henning Kunz

## Abstract

Bacteria can trigger protein mistranslation to survive stress conditions ^1^. Mitochondria and plastids evolved from bacteria and therefore also use prokaryotic-type expression machineries to synthesize proteins. However, fungi and animal mitochondria are highly sensitive to mistranslation, which for instance manifests in lethal mitochondrial cardiomyopathy disorder ^2^. The response in plant cells is unknown. Glutaminyl-transferRNAs (Gln-tRNA^Gln^) of bacteria, mitochondria, and plastids are synthesized indirectly ^3,4^. Initially, tRNA^Gln^ gets charged with glutamate. Subsequently, Gln is produced through trans-amidation by the aminoacyl-tRNA amido-transferase complex GatCAB. Consequentially, affected GatCAB activity results in pools of misloaded Glu-tRNA^Gln^. Here we show that Arabidopsis mutants with decreased GatCAB level provide global insights into organellar mistranslation in plants. Proteomics revealed Gln-to-Glu misincorporation in plastid- and mitochondrially-expressed protein complexes with only modest abundance changes in mutant plants. Plastids appear more lenient to mistranslation as they exhibit much higher Gln-to-Glu misincorporation. Through efficient compensatory mechanisms, mutant plants display surprisingly subtle phenotypes. However, their acclimation to temperature stress differs. Interestingly, wild-type plants under similar stress also have altered Gln-to-Glu misincorporation. Our study shows that the response towards organellar mistranslation varies among eukaryotes. In plants, this knowledge can be used to improve stress tolerance.

## Main

In the central dogma of molecular biology, Crick described the directionality of information transfer in cells (DNA → RNA → protein) ^5,6^. Later he clarified, the dogma says “nothing about errors” and added: “It was assumed that, in general, the accuracy of transfer was high.” ^7^. This was confirmed for DNA replication and transcription. In contrast, ribosomal mis-decoding or transfer-RNA (tRNA) mis-acylation render protein synthesis an error-prone step (amino acid misincorporation 10^−3^ to 10^−4^ per codon) ^8–10^. A source for tRNA mis-acylation in bacteria is the GatCAB-dependent pathway to produce Gln-tRNA^Gln^ from Glu-tRNA^Gln^ (Fig. 1a). A similar pathway exists less frequently for the transamidation of Asp-tRNA^Aln^ to Aln-tRNA^Aln^ ^11^. Disturbed GatCAB activities yield mis-acylated tRNAs resulting in amino acid misincorporation ^1,12^. Mistranslation seems unfavorable, newly made proteins can be unfunctional and the energy investment for protein synthesis is lost ^13^. However, bacteria are surprisingly tolerant to mistranslation and under stress conditions even trigger it ^11^. Many Rifampicin-resistant clinical Mycobacteria strains alter GatCAB activity, which yields more amino acid misincorporations. The resulting proteome plasticity is beneficial under adverse growth conditions, such as antibiotic selection pressure, since mistranslated RNA-polymerase variants no longer bind Rifampicin and remain functional ^12^ ^14^. Hence, prokaryotes may have kept the indirect Gln/Asp-synthesis pathways throughout evolution to diversify their proteome without changing the highly error-protected DNA code ^11^.

**Fig. 1:**
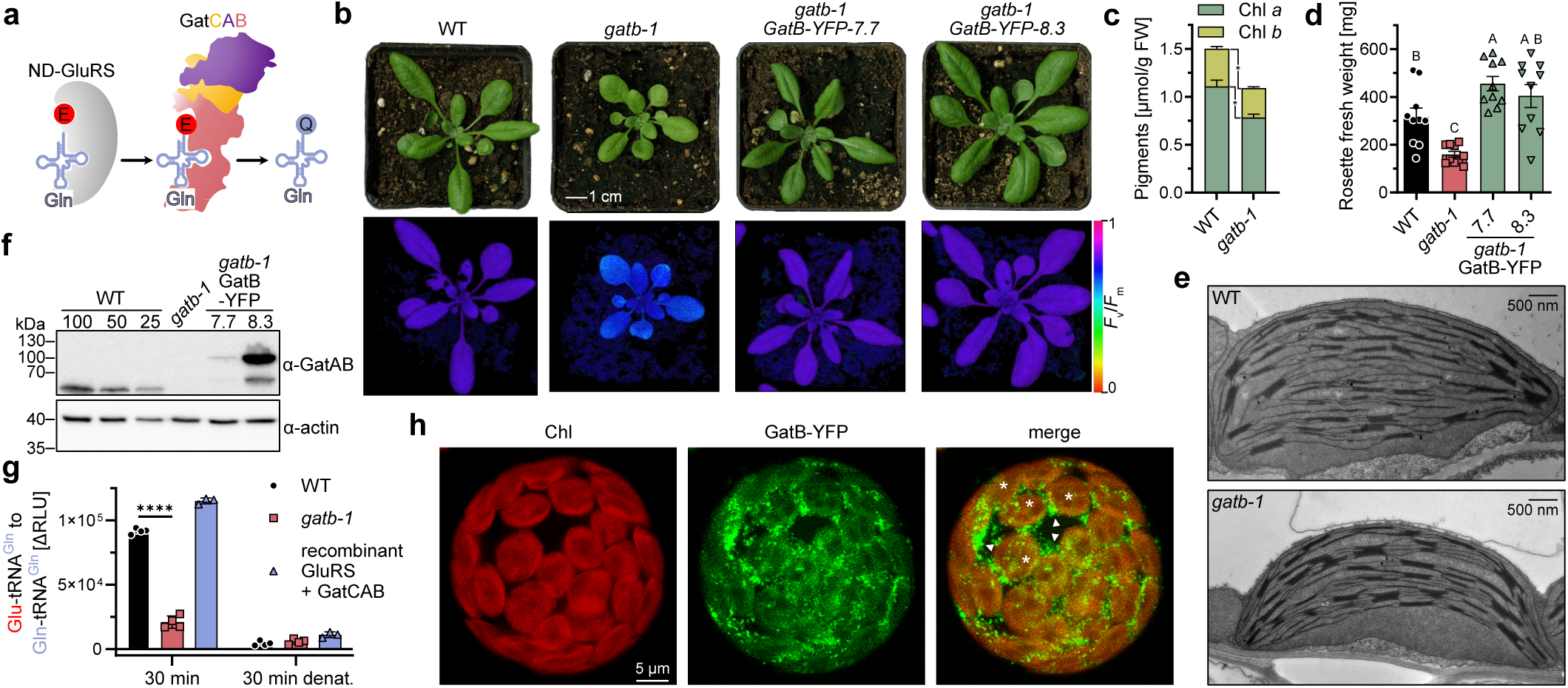
A hypo-morphic mutant of mitochondria and chloroplast stroma localized *GatB* shows decreased enzymatic GatCAB activity and subtle phenotypes. (a) Cartoon representation of the GatCAB mediated trans-amidation of Glu-tRNA^Gln^ to Glu-tRNA^Gln^. (b) Photographs and PAM images (*F*_v_/*F*_m_) of WT, *gatb-1*, *gatb-1 ProGatB:GatB-YFP*-*7.7* and *-8.3* plants. (c) Chl *a* (dark green) and Chl *b* (yellow green) in WT and *gatb-1* plants. Mean +/- SEM; n=3; Multiple unpaired t-tests with correction for multiple testing according to Benjamini, Krieger, and Yekutieli using a false discovery rate (FDR) of 5% indicated by the asterisks. (d) Quantification of above ground plant fresh weight (in mg) of WT (black), *gatb-1* (red), *gatb-1 ProGatB:GatB-YFP*-*7.7* and -*8.3* (green) plants. Mean +/- SEM; n=10; Ordinary one-way ANOVA. Letters indicate significantly different groups (p value ≤ 0.05). (e) Transmission-electron microscopy of WT and *gatb-1* chloroplasts. (f) Immunoblot detection of GatAB (molecular weights including transit peptides: GatB = 60.9 kDa, GatB-YFP = 88.9 kDa and GatA = 75.2 kDa) levels in normalized total leaf extracts of WT, *gatb- 1* and two independent complementation lines (*gatb-1 ProGatB:GatB-YFP-7.7* and *-8.3*). GatA and GatB proteins run approximately at the same size ^3^. Different amounts of WT extract (100%, 50%, and 25%) were loaded for easier comparison. The α-actin antibody served as loading control. See also Extended Data Fig. 1i. (g) Label-free quantitative biochemical assay probing the trans-amidation activity of WT (black) and *gatb-1* (red) stromas as well as recombinant ND-GluRS + GatCAB proteins (blue). Mean +/- SEM; n = 4; Two-way ANOVA with Tukey correction for multiple comparisons. Only significantly different *gatb-1* vs. WT comparisons (at 30 min and 30 min denatured) are indicated in the graph. ****: adj. p-value ≤ 0.0001. A detailed description, purification of recombinant proteins, loading controls and raw data of the assays can be found in Extended Data Fig. 3. (h) Confocal laser microscopy of *ProUBQ10:GatB-YFP* Arabidopsis protoplasts. Chlorophyll auto- fluorescence in red (left), GatB-YFP fluorescence in green (middle) and a merged image (right). Arrowheads indicate mitochondria while asterisks mark chloroplasts.

Endosymbiotic organelles in eukaryotic cells also rely on the indirect Gln-tRNA^Gln^ pathway to synthesize essential proteins encoded on their bacterial genomes. The reasons for this dependency are even more obscure. Studies on yeast, animal, and human mitochondria have shown that many mutations in GatCAB subunits disrupt oxidative phosphorylation and are lethal ^2,15–18^. Silencing *GatA* in mice cells arrests mitochondrial protein synthesis without amino-acid misincorporation because the translational apparatus rejects mis-acylated tRNA^Gln^ ^17^. This suggests that mitochondria generally do not tolerate mistranslation. Plant cells are more complex harboring a second bacterial-derived organelle, the chloroplasts. Photosynthesis depends on protein synthesis within chloroplasts and also hinges on intact metabolism in mitochondria ^19^. The robustness of plant cells towards organellar mistranslation is unknown.

### GatCAB is essential but decreased activity well tolerated

To explore a plant cell’s tolerance to organellar mistranslation, we attempted to isolate *Arabidopsis thaliana* (Arabidopsis) mutants devoid of GatCAB subunits (Fig. 1a). Three Arabidopsis lines carrying T(ransfer)-DNA insertions in *GatA (gata-1)*, *GatB (gatb-2),* and *GatC* (*gatc-1*) were exclusively found as heterozygotes and never yielded homozygous mutant progenies (Extended Data Fig.1a-g) indicating that GatCAB is vital for seedling establishment. However, we isolated homozygous *gatb-1* mutants carrying a T-DNA insertion 118 bps upstream of the start codon (Extended Data Fig.1c). *gatb-1* plants exhibit only mild phenotypic variations from wild type (WT) i.e., slightly decreased maximum efficiency of photosystem II (PSII) (*F*_v_/*F*_m_) (Fig. 1b), somewhat paler leaves (∼30% decreased total chlorophyll content; Fig. 1c) and slower growth (∼55% decreased fresh weight, Fig. 1d). Chloroplast morphology in *gatb-1* resembled WT with intact envelope membranes and thylakoids characteristically separated into stroma lamella and grana stacks (Fig. 1e).

Whole-genome sequencing of backcrossed *gatb-1* plants confirmed that observed phenotypes are linked to the 5’-upstream insertion in the *GatB* locus; no other homozygous second-site insertion/mutation were present (Extended Data Fig. 1c; see also methods). Reintroducing GATB-YFP into homozygous *gatb-1* plants recapitulated WT appearance (Fig. 1b,d). *gatb-1* was also crossed to heterozygous *gatb-2/GATB* (Extended Data Fig. 2a). As expected, 50% of F1 plants (*gatb-1*/*gatb-2* genotypes) displayed similar but exacerbated phenotypes as *gatb-1* (Extended Data Fig. 2 b,c), which was also documented in *GatA* post-transcriptionally silenced lines (*amiGatA*, Extended Data Fig. 2d,e).

In immunoblots using an antibody that detects GatA and GatB (Extended Data Fig. 1i and 3), both subunits were undetectable in *gatb-1* or *amiGatA* (Fig. 1f, and Extended Data Fig. 1i, 2f) suggesting that GatCAB subunits are co-regulated and residual amounts in the mutants are low. Equal amounts of WT and *gatb-1* stromal total protein (Extended Data Fig. 3e) revealed ∼70% decreased Glu-tRNA^Gln^ to Gln- tRNA^Gln^ conversion in *gatb-1* (Fig. 1g and Extended Data Fig. 3a,f for raw values and all controls of the established label free trans-amidation assay). The GatB-YFP protein localized to mitochondria and chloroplasts (see merge with Chl fluorescence Fig. 1h). Dual-localization of GatB is in line with a previous study ^3^. For plastid suborganellar studies, WT chloroplasts were fractionated and immunoblotted. The results showed that chloroplast GatCAB complexes reside in the stroma (Extended Data Fig. 1j).

In summary, GatCAB activity is essential for plant reproduction. However, a strong reduction, which results in decreased Glu-tRNA^Gln^ trans-amidation, is surprisingly well tolerated by the plant.

### Reduced GatCAB activity triggers Q→E mistranslation in plastid and mitochondrially-expressed proteins

In prokaryotes, reduced GatCAB activity increases Gln(Q) to Glu(E) (Q→E) and/or Asn(N) to Asp(D) (N→D) amino acid misincorporation ^12,14^. However, for organelle-synthesized proteins in eukaryotes this has not been shown ^17,20^.

Using shotgun proteomics, we compared the percentage of unique peptides with either Q→E or N→D amino acid misincorporations in isolated mitochondria and chloroplasts from all genotypes. We found no striking differences in the percentage of unique peptides with either Q→E or N→D amino acid misincorporation for proteins expressed in the cytosol (CYTex) or peptides with N→D substitutions stemming from plastid (PLex) or mitochondrial (MTex) protein synthesis (Fig. 2a,b). However, *gatb-1* mutants revealed a clear increase of peptides with Q→E substitutions in MTex and PLex peptides, contrasting WT organelles where no such misincorporation was detected in PLex and only a low number in MTex (Fig. 2a,b).

**Fig. 2:**
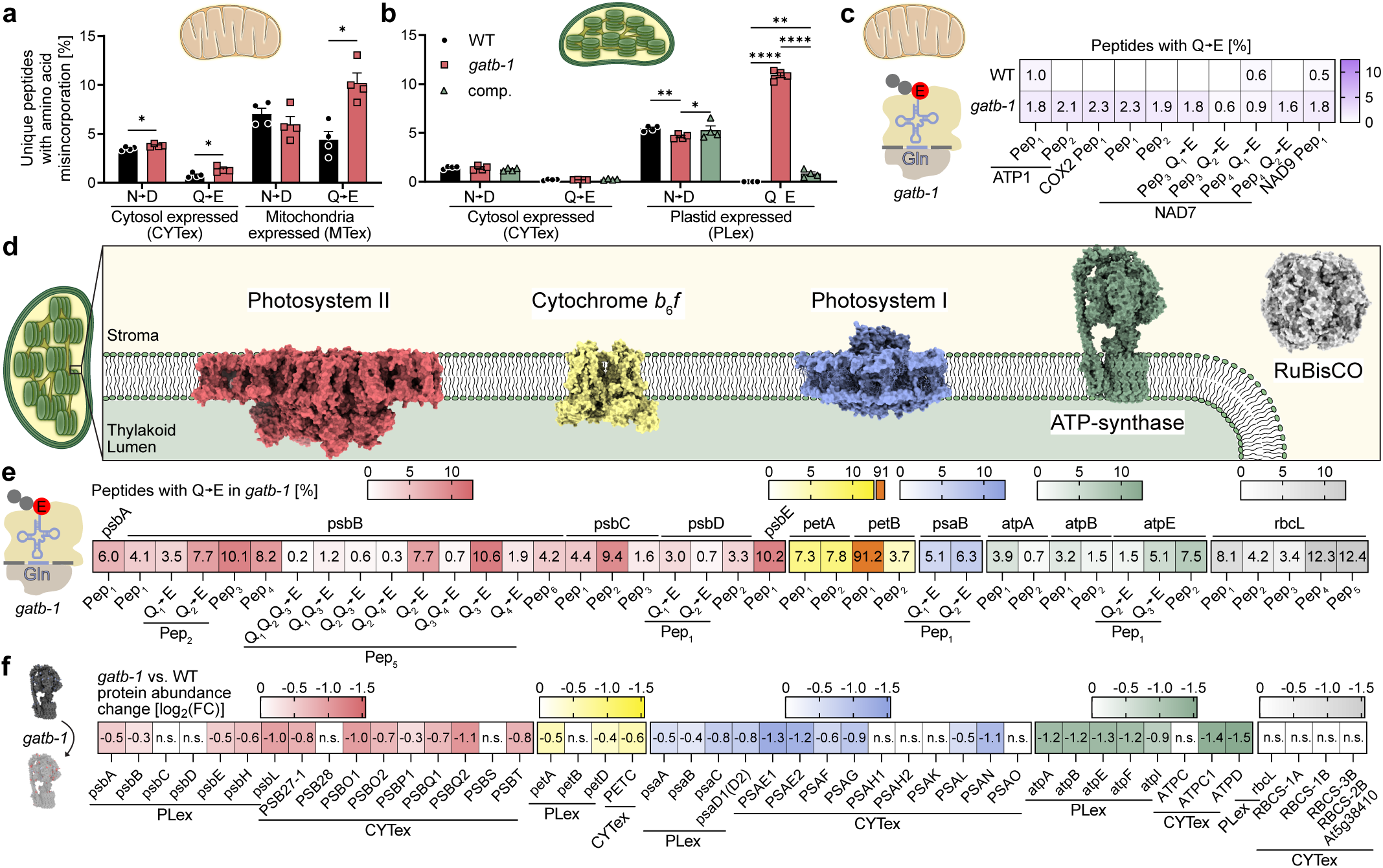
Reduced GatB or GatA results in increased Q→E mistranslation in plastid and mitochondrially expressed proteins and perturbed chloroplast protein homeostasis. (a) Percentages of unique peptide with N→D or Q→E substitutions in CYTex or MTex proteins in WT (black) or *gatb-1* (red) measured by mitochondria proteomics. Mean +/- SEM; n= 4; Multiple unpaired t-tests with multiple testing correction according to Benjamini, Krieger, and Yekutieli using a false discovery rate (FDR) of 5%. A single asterisk indicates a significant difference with the indicated FDR. (b) Percentages of unique peptides with either N→D or Q→E substitutions in either CYTex or PLex proteins in WT (black), *gatb-1* (red) or the *gatb-1* complementation (comp.; green bars) assessed by chloroplast proteomics. Mean +/- SEM; n= 4; Two-way ANOVA with Tukey correction for multiple comparisons. * = adj. p-value ≤ 0.05, ** = adj. p-value ≤ 0.01, **** = adj. p-value ≤ 0.0001. (c) Q→E misincorporation rates of individual MTex peptides in WT and *gatb-1* measured by mitochondria proteomics. Mean values are shown; n= 3-4; (d) Surface representation of major multi-subunit protein complexes involved in photosynthesis in plant chloroplasts: RUBISCO (grey; PDB-ID:4hhh), ATP-syn (green; PDB-ID:6fkf), photosystem I (PSI; blue; PDB-ID:5l8r), Cytochrome *b*_6_*f* (yellow; PDB-ID: 6rqf) and photosystem II (PSII; red; PDB-ID:3jqu). (e) Q→E misincorporation rates of individual peptides of PLex proteins in *gatb-1* measured by chloroplast proteomics. Color code: RuBisCO (grey), ATP-syn (green), photosystem I (PSI, blue), Cytochrome *b*_6_*f* (yellow) and photosystem II (PSII; red). Mean values are shown; n= 3-4. (f) Protein abundance changes (fold change; FC; *gatb-1* vs. WT) in subunits of protein complexes listed in (c) determined by label-free chloroplast proteomics. Color code: RuBisCO (grey), ATP-syn (green), photosystem I (PSI, blue), Cytochrome *b*_6_*f* (yellow) and photosystem II (PSII; red). Significance was determined by t-tests in combination with a permutation based false-discovery rate (FDR) of 5% and s0=0.1.

In mitochondria, we identified 81 unique peptides (from 17 MTex protein groups irrespective of their modifications) in ≥ 3 biological replicates, 33 of which have glutamines. Ten individual Q→E changes were identified in eight different peptides from ATP synthase subunit1 (ATP1), cytochrome c oxidase 2 (COX2), and NADH-dehydrogenase subunits (NAD) 7 and 9 (Fig. 2c). The mean misincorporation rate was 1.6% in *gatb-1*. Individual rates never surpassed 2.1%. We also found three peptides from WT mitochondria carrying Q→E misincorporations (all below 1% rate), one in ATP1 and one each in two NAD subunits (Fig. 2c).

In *gatb-1* chloroplasts, we identified 321 distinct peptides (irrespective of their modifications) from 58 uniquely plastome-encoded protein groups in ≥ 3 biological replicates. Out of these, 142 peptides of 48 protein groups contain glutamines (Supplementary Table 2). 21.1% of the Q-containing peptides in 10.4% PLex proteins carried one or multiple Q→E substitutions in *gatb-1*. Not a single peptide was found in WT. Q→E misincorporations in *gatb-1* were present in peptides known to assemble into major plastid protein complexes i.e., photosystem II (PSII), cytochrome *b*_6_*f* (Cyt*b*_6_*f*), photosystem I (PSI), ATP-synthase (ATP-syn) and Ribulose-1,5-bisphosphat-carboxylase/-oxygenase (RuBisCO) (Fig. 2d,e). Inherent to the experimental restrictions of mass spectrometry, we cannot interpret missing peptides/proteins. However, we detected 16 Q-containing peptides of seven plastid-expressed subunits of the NAD(P)H-dehydrogenase (NDH) complex and 24 Q-containing peptides in 13 plastid-expressed subunits of the 70S ribosome. Notably, no Q→E-altered peptides in these complexes were found (Supplementary Table 1).

The rates for single Q→E misincorporation varied greatly among PLex peptides with mean percentages ranging from 0.72% (atpA Pep2) to 91.2% (petB Pep1; Fig. 3e). The mean Q→E misincorporation rate for all PLex peptides detected in *gatb-1* was 8.5%. psbB Pep5 in the PSII reaction center (Extended Data Fig. 4a), which contains three Qs, revealed single, double, and triple Q→E substitutions in different combinations and percentages (Fig. 3e and Extended Data Fig. 4b). Strikingly, petB Pep1 of the H^+^ pumping Cyt*b*_6_*f* complex showed a 91.2% Q→E misincorporation rate in *gatb-1* (Fig. 2e). The substituted Q localizes near the luminal exit of the H^+^ conducting pore, less than 20 Å away from the electron-carrying plastoquinone ring, which may alter its coordination in mutant Cyt*b*_6_*f* complexes (Extended Data Fig. 4c).

**Fig. 3:**
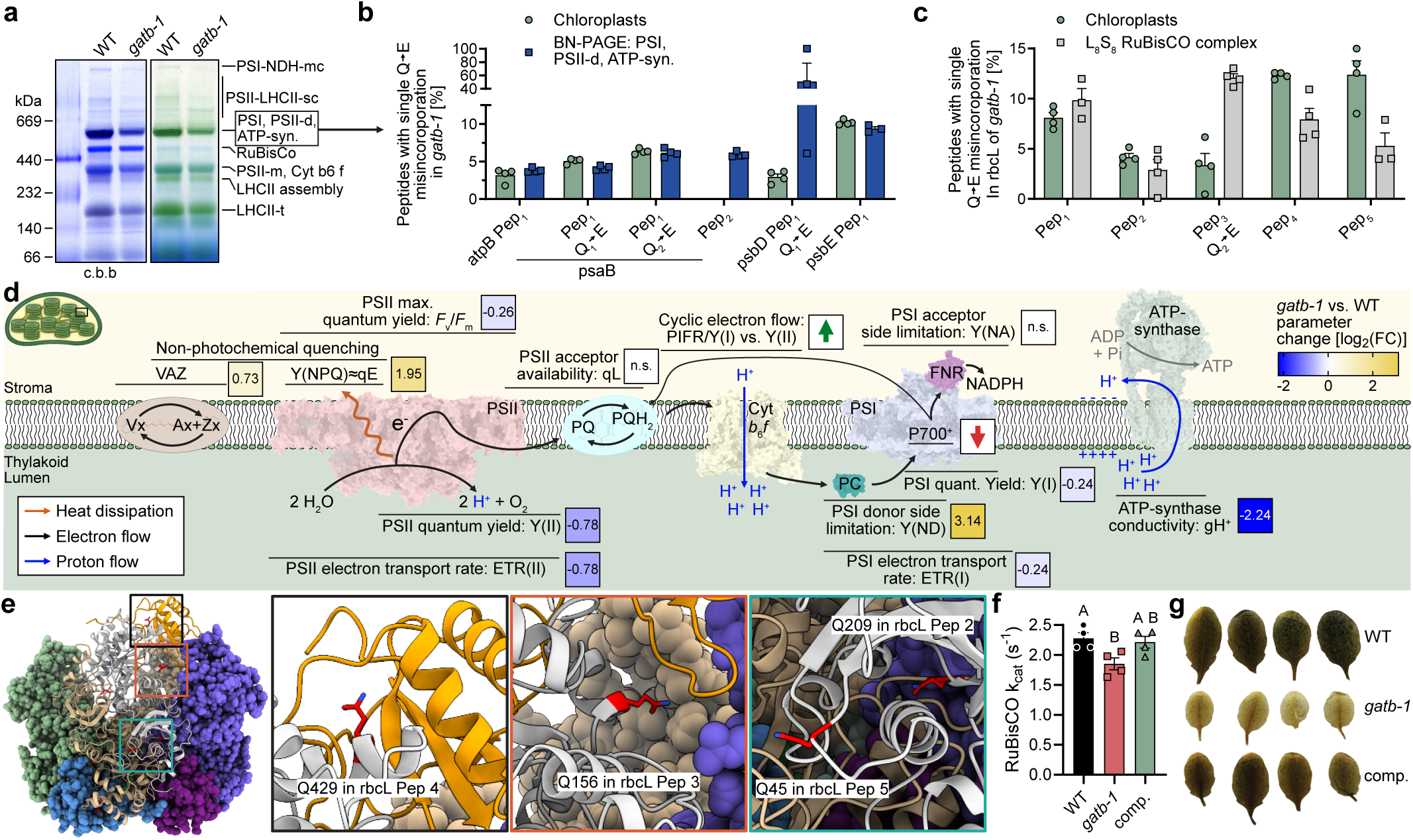
*gatb-1* plants exhibit mild yet distinct changes in the assembly of photosynthetic complexes, photosynthetic parameters, RuBisCO activity and starch accumulation. (a) BN-PAGE analysis photosynthetic complexes from WT and *gatb-1* chloroplasts. The same area of the gel is depicted before (right) and after (left) coomassie staining. The box indicates the band used for the proteomics experiment in (b). For more biological replicates please refer to Extended Data Fig. 8a. mc: mega complex; sc: supercomplex; m: monomer; t: trimer. (b) Q→E misincorporation rates of assembled protein complexes determined by proteomics on a cut out PSI, PSII-d and ATP-syn band of a BN-PAGE (blue) and chloroplasts (green, already shown in Fig. 2e) in *gatb-1*. Mean +/- SEM; n= 3-4. (c) Q→E misincorporation rates in the RuBisCO large subunit (rbcL) of isolated chloroplasts (green; already shown in Fig. 2e) and combined L_8_S_8_ RuBisCO containing SEC fractions (gray) in *gatb-*Mean +/- SEM; n= 3-4. (d) Schematic and simplified overview of processes during photosynthesis and parameters measured in this study. Significant parameter changes (fold change; FC) measured at 53 PAR in *gatb-1* vs. WT are shown. n.s.= not significant. Multiple t-tests in combination with a permutation-based false-discovery rate (FDR) of 5% and s0=0.1. Increased PIFR/Y(I) vs.Y(II) and decreased P700^+^ in *gatb-1* vs. WT are indicated by a green or red arrow, respectively. Detailed graphs for all parameters can also be found in Extended Data Fig. 8. (e) Mapping of mistranslated Q residues onto the structure of the RuBisCO complex. Arabidopsis large and small subunits (PDB ID: 5iu0) were superimposed onto the L_8_S_8_ pea RuBisCO (PDB IDs: 4mkv). Large subunits are colored in light gray and sand colors while the small subunit is depicted in orange. Mistranslated Q residues which are resolved in the structure are colored in red (Q45, Q156, Q209 and Q429). The regions of the peptides with misincorporations are zoomed in (black, turquoise and orange boxes). (f) Per molecule RuBisCO reaction rates (*kcat*) of WT (black), *gatb-1* (red) and a complementation line (comp.; green). Mean +/- SEM; n=4; Ordinary one-way ANOVA. Letters indicate significantly different groups. (g) Transitory starch stain with Lugol’s iodine in WT, *gatb-1* and comp. leaves after 4 hours of light.

We also analyzed the leaf proteome of phenotypically more severe *amiGatA* mutants. Although, the normalized number of Q→E peptides was similar between *gatb-1* and *amiGatA*, the abundance of individual Q→E peptides in ≥ 3 biological replicates and mean misincorporation rates were mostly higher in *amiGatA* (Extended Data Fig. 5a-c; mean mis-incorporation rate in *gatb-1* = 5.8% and in *amiGatA* = 9.4%). No PLex peptides with Q→E were identified in WT. Hence, Q→E misincorporation rates may scale with phenotypic severity.

By crossing *gatb-1* with a trans-plastomic YFP (plYFP) reporter line ^21^, we tested if plastome-encoded transgenes become mistranslated. YFP contains several Q residues sensitive to Glu replacements (Q69 and Q94; ^22,23^). Resulting WT-plYFP and *gatb-1-plYFP* progenies expressed similar YFP levels (Extended Data Fig. 6a-c). However, YFP fluorescence in *gatb-1-plYFP* was 36% below WT-plYFP (Extended Data Fig. 6d). A 17% Q→E rate in the YFP protein (no Q→E found in WT-*plYFP*; Extended Data Fig. 6f-h) confirmed that the fluorescence loss likely originates from Q→E substitutions unique to *gatb-1-plYFP* plants.

In summary, altering GatCAB levels resulted in strongly increased Q→E misincorporation of organelle- expressed proteins with more dramatic effects in plastids.

### Lack of GatCAB leads to perturbed chloroplast protein homeostasis

As a consequence of amino acid mistranslation, protein misfolding and degradation are expected ^8^. To probe protein abundances, we performed label-free quantitative (LFQ) proteomics on isolated chloroplasts (Fig. 2f). *gatb-1* revealed 171 significantly changed proteins in the chloroplast proteomics dataset while protein abundances in complemented mutants resembled WT (Extended Data Fig. 7a,b and Supplementary Table 2). As exemplified for PSII, Cyt*b*_6_*f*, PSI, ATP-synthase (ATP-syn), and RuBisCO, it appears that protein complexes respond differently to the stress (Fig. 2e,f). For instance, despite an 8.1% mean Q→E misincorporation rate in the large RuBisCO subunit, neither the abundance of the plastome-encoded rbcL nor the nuclear-encoded RbcS subunits were significantly altered. Conversely, almost all detected ATP-syn components, with below average Q→E misincorporation rates, were clearly decreased (up to -1.5 log2FC). Notably, petB, the Cyt*b*_6_*f* subunit containing a peptide with 91.2% Q→E, was not significantly altered. Other detected Cyt*b*_6_*f* subunits with Q→E changes were only mildly reduced in *gatb-1*. PSII and PSI both showed varying degrees of lower subunit abundances with nuclear-encoded subunits being more strongly decreased than plastome-encoded ones. Two of which, psbC and psbD, were not significantly decreased although both carried Q→E misincorporations. Also, the psbB PSII subunit, with the aforementioned Q-rich Pep5 (overall 25% Q→E misincorporations), was barely impacted (Fig. 2e,f). Plastid protein abundance changes were confirmed by representative immunoblots (Extended Data Fig. 7c).

Several general conclusions can be drawn: 1) all Q→E-modified PLex proteins in *gatb-1* were unchanged or significantly decreased compared to WT; 2) not only plastome-encoded but also nuclear-encoded subunits from the same complexes accumulated at lower abundance during stromal Q→E misincorporation stress; 3) proteins outside aforementioned mistranslated complexes i.e., without Q→E can either be up or downregulated in response to mistranslation stress; 4) protein abundances of mitochondria-encoded (MTex) proteins were not significantly altered in *gatb-1* (Extended Data Fig. 7d). 5) LFQ analysis of more affected *amiGatA* plants showed similar but more drastic protein abundance changes than in WT and *gatb-1* (Extended Data Fig. 5d-g, Supplementary Table 3) indicating that, similar to Q→E rates, protein abundance changes follow phenotypical severity.

Since Q→E misincorporations are much more prevalent, have stronger effects on protein abundance in plastids, and their physiological response has not been explored, we mainly focused on *gatb-1* chloroplasts for the remaining study.

### Mistranslated peptides in protein complexes alter photosynthesis

To exclude that Q→E mistranslated proteins undergo degradation and never assemble into multi-subunit complexes, we probed oligomeric protein states. Fully-assembled light reaction machinery complexes from WT and *gatb-1* plastids were separated by native PAGE. In line with LFQ proteomics (Fig. 2f), band inspection revealed intensity differences in PSII-light harvesting complex II (LHCII) super complexes (PSII-LHCII-sc) and PSI-NDH megacomplex (PSI-NDH-mc) formation. Also, LHCII trimers (LHCII-t), LHCII assembly, PSII monomer (PSII-m), Cyt*b*_6_*f*, and PSI, PSII dimer (PSII-d) and ATP-syn were slightly less abundant in *gatb-1*. Notably, no missing complexes were observed in *gatb-1* suggesting that assembly of Q→E misincorporated subunits into protein complexes is possible (Fig. 3a and Extended Data Fig. 8a). To probe this assumption, a high molecular weight band (PSI, PSII-d, ATP- syn) and the fully-assembled L_8_S_8_ hexadecamer RuBisCO complex from WT and *gatb-1* were isolated (Extended Data Fig. 8a and 9a,b) and subjected to mass spectrometry. For ATP-syn, PSI, PSII, and rbcL overall similar, sometimes even higher, Q→E misincorporation rates were found in isolated complexes as in *gatb-1* chloroplasts extracts (Fig. 3b,c). This confirms that peptides with Q→E substitutions are present in fully-assembled multi-subunit protein complexes.

To understand how *gatb-1* performs photosynthesis while challenged with partly mistranslated protein complexes several PSII and PSI-related parameters were assayed (Fig. 3d). As mentioned before, maximum PSII efficiency (*F*_v_/*F*_m_) in WT was at 0.77 while *gatb-1* scored 0.64 indicative of moderate PSII damage (Fig. 3d). Most genotype-specific parameters differed in low to medium light intensities (Extended Data Fig. 8b-8j). Under these conditions, incoming light energy was differently distributed among genotypes (Fig. 3d and Extended Data Fig. 8k-aa): non-photochemical quenching (Y(NPQ) ≈ energy-dependent quenching (qE)) in *gatb-1* was above WT whereas non-regulated energy dissipation Y(NO) was unchanged, and Y(II) (PSII quantum yield) markedly decreased in the mutant. PSII acceptor availability (qL) was similar. Linear electron transport rate ETR (II) was lower in *gatb-1,* pointing at a blockade in Cyt*b*_6_*f* that led to more PSI donor side limitation (Y(ND)). PSI efficiency was only mildly affected in mutant plants (Y(I)) without PSI acceptor side limitation (Y(NA)). The slow P700 oxidation kinetics in *gatb-1* during far-red light exposure (P700^+^), is in line with the observed higher Y(ND) ^24^. gH^+^, a proxy for ATP synthase activity ^25^, was a fraction (20%) of the WT value in *gatb-1* owing to the low abundance of ATP-syn complexes (Fig. 2f and Extended Data Fig. 7c). In addition, we documented indications for high cyclic electron transport (CET) in *gatb-1* (increased dark-induced fluorescence rise (PIFR) and increased slope Y(I) vs Y(II) ^26,27^). Q→E misincorporations in *gatb-1* photosystems may also affect movement of LHCII from PSII to PSI via STN7-catalyzed LHCII phosphorylation (qT) ^28^, since *gatb-1* by trend had lower qT than WT. Lastly, carotenoids which contribute to qE and qZ ^29^, a third important NPQ component, were quantified. While Violaxanthin (vx) was unaltered, de-epoxidated molecules antheraxanthin (Ax) and zeaxanthin (Zx) accumulated in *gatb-1* (sum of Vx, Ax, Zx = VAZ). The VAZ precursor β-carotene (Car) was decreased whereas the α-xanthophyll lutein, a potent quencher of triplet chlorophyll, increased in *gatb-1* (Fig. 3d and Extended Data Fig. 8k-aa).

In summary, we found that Q→E misincorporations persist in fully-assembled complexes of *gatb-1*’s photosynthetic machinery. PSII function is only mildly affected but linear electron flow towards PSI is decreased. *gatb-1* utilizes several NPQ components (qE, qZ, qT) to safely release surplus energy successfully maintaining sufficient photosynthesis and growth.

### Q→E misincorporations affect CO2-fixation

To probe the effect of Q→E changes on protein function, RuBisCO represents a premier target: by quantifying the number of active sites via stoichiometric binding of radiolabeled [^14^C]carboxy-arabinitol bisphosphate (^14^CABP) ^30^ it is possible to assay enzyme activity per molecule RuBisCO.

Q→E misincorporated peptides mapped to the rbcL surface and near the catalytic center of the fully-assembled L_8_S_8_ RuBisCO hexadecamer (Fig. 3e). Interestingly, the CO2-fixation rate of RuBisCO (*k*cat) dropped significantly by 23% in *gatb-1* compared to WT and complemented mutants (Fig. 3f). In line with this result, we found less transitory leaf starch, the main CO2 storage during the day, in *gatb-1* than in control plants (Fig. 4g).

**Fig. 4:**
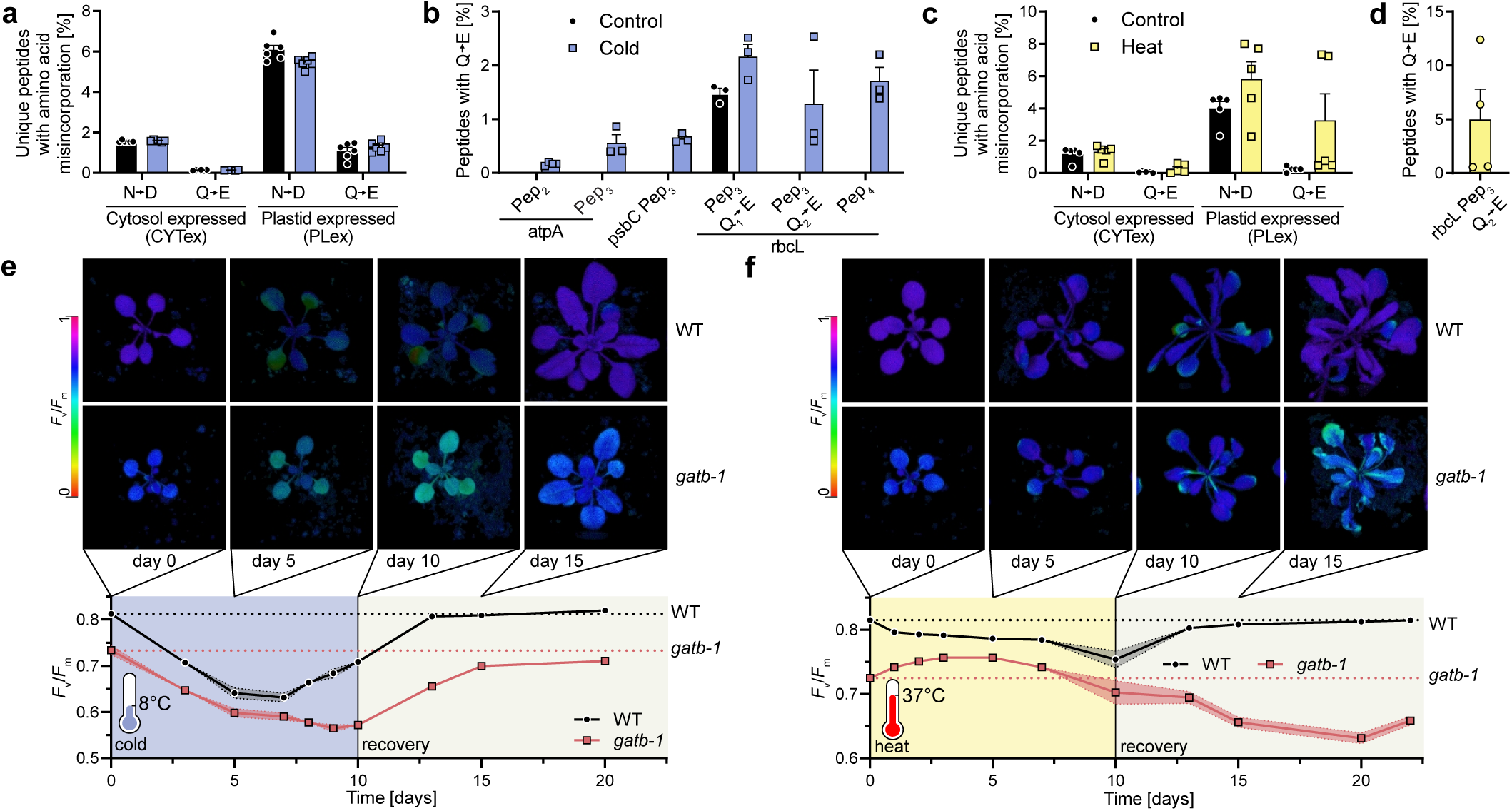
Plastid Q→E misincorporation rates are growth temperature-dependent, which affects the acclimation capacity of *gatb-1* mutants. (a and c) Percentage of unique peptides with either N→D or Q→E substitutions of WT either subjected to cold (blue, a) or heat (yellow, c) treatment in CYTex or PLex proteins measured by leaf proteomics (data from^34^). Unique mistranslated peptides were normalized with the total number of unique peptides. Mean +/- SEM; n= 5-6. (b and d) Q→E misincorporation rates of peptides from WT plants either subjected to cold (blue, a) or heat (yellow, d) treatment (in%) measured by leaf proteomics (data from^34^). Mean +/- SEM; n= 3-5. (e and f) Maximum PSII efficiency (*F*_v_/*F*_m_) in WT (upper row and black circles) or *gatb-1* (lower row and red squares) plants treated with cold (8°C; blue; e) or heat (37°C; yellow; f) for 10 days followed by recovery at ambient temperatures (22°C). Mean +/- SEM; n= 8-10. All WT and *gatb- 1* values were significantly different for each measured data point tested with multiple unpaired t-tests with correction for multiple testing according to Benjamini, Krieger, and Yekutieli using a false discovery rate (FDR) of 5%.

In essence, our findings from RuBisCO assays show that tolerated Q→E misincorporations in *gatb-1* can alter protein function.

### Plastid Q→E misincorporation is temperature-dependent, affecting plant acclimation capacities

Mistranslating bacteria exhibit increased heat tolerance ^31^. Similar scenarios may exist in endosymbiotic organelles. At 10°C versus 30°C plastids synthetize twice as much RuBisCO to compensate the enzymatic activity loss ^32^, whereas heat triggers RuBisCO turnover to outfit L_8_S_8_ oligomers with the temperature-stable RbcS-B subunits ^33^. It follows that heat and cold stress may affect amino acid misincorporation in plastids. We tested this by reanalyzing proteomics data from temperature-shifted plants i.e., 20°C to 15°C or 25°C ^34^. We focused on N→D and Q→E changes. In the cold, the overall number of peptides with N→D or Q→E was unchanged. However, upon closer inspection six PLex peptides, two of the ATP-syn, one in PSII and three from rbcL showed unique or higher Q→E misincorporation rates in cold-treated wild-type plants (Fig. 4a,b). In heat-stressed plants, trends for higher N→D and Q→E misincorporation were seen only for PLex peptides. Again, we found a rbcL peptide (Pep2), which carried Q→E changes exclusively in 37°C-treated wild-type plants (Fig. 4c,d).

Since translation accuracy seems to vary in WT plants grown at different temperatures, we tested the acclimation capacity of the Q→E hyper-mistranslating mutant *gatb-1*. 20 days-old plants, raised at ambient conditions (22°C), were shifted either to 8°C or to 37°C for 10 days followed by a 10-days recovery phase at 22°C. Chlorophyll fluorescence was recorded as a sensitive non-invasive reporter for the plant’s temperature tolerance ^35,36^.

Confirming previous studies ^37^, cold shifts resulted in a seven-day-long *F*_v_/*F*_m_ decline in all genotypes (Fig. 4e). Interestingly, at the seven-day mark WT successfully acclimated to low temperatures i.e., *F*_v_/*F*_m_ increased close to the starting value. In contrast, *gatb-1* plants did not show *F*_v_/*F*_m_ recovery if cold conditions persisted. Only a reset to ambient growth temperatures triggered *F*_v_/*F*_m_ increase in *gatb-1* plants. Nevertheless, the mutants’ acclimation was slower than in wild-type plants and, distinct from WT, never reached pre-cold-treatment *F*_v_/*F*_m_ levels (Fig. 4e).

Heat treatments gave a different acclimation pattern. In WT plants *F*_v_/*F*_m_ decreased much less as in the cold (Fig. 4f). After day seven days at 37°C, this decay seemed to accelerate. Conversely, *gatb-1* plants displayed an initial *F*_v_/*F*_m_ gain that faded after day seven. Yet another trend emerged when plants returned to ambient temperatures: While WT rapidly reset *F*_v_/*F*_m_ to pre-treatment level, the mistranslating *gatb-1* mutants lost *F*_v_/*F*_m_ until finally, 20 days after the beginning of the treatment, *F*_v_/*F*_m_ recovered (Fig. 4f).

### Plants device compensatory mistranslation mechanisms

Overall, temperature stress induced numbers and rates of Q→E peptides are lower in WT than in Q→E hyper-mistranslating *gatb-1* plants, which display surprisingly mild phenotypes. This prompted us to identify the highly effective compensatory mechanisms of endosymbiotic organelles in response to Q→E stress. To that end we quantified mRNA transcripts in all genotypes. 1461 differentially expressed genes (DEGs) in *gatb-1* vs WT (663 up- and 798 down-regulated) were identified (Fig. 5a and Supplementary Table 4), whereas transcription profiles in complemented mutant lines were close to WT (Extended Data Fig. 10a). Biological functions of upregulated DEGs were assigned by gene ontology (GO) enrichments. Most upregulated genes fell into GO categories summarized as “chloroplast and mitochondrial protein synthesis and biogenesis”. This included GO terms plastid organization, plastid transcription, protein re-folding, mitochondrion organization, mitochondrial gene expression and ribosome biogenesis. The second cluster of upregulated GO terms can be coined “metabolic responses” (Fig. 5b, Extended Data Fig. 10b,c for interesting individual genes within these GO terms; see Supplementary Table 5 for all enriched GO terms). Given the importance of chloroplast and mitochondria for metabolism these responses seem partly consequences of malfunctions within the first GO category affected in *gatb-1* (“chloroplast and mitochondrial protein synthesis and biogenesis”).

**Fig. 5:**
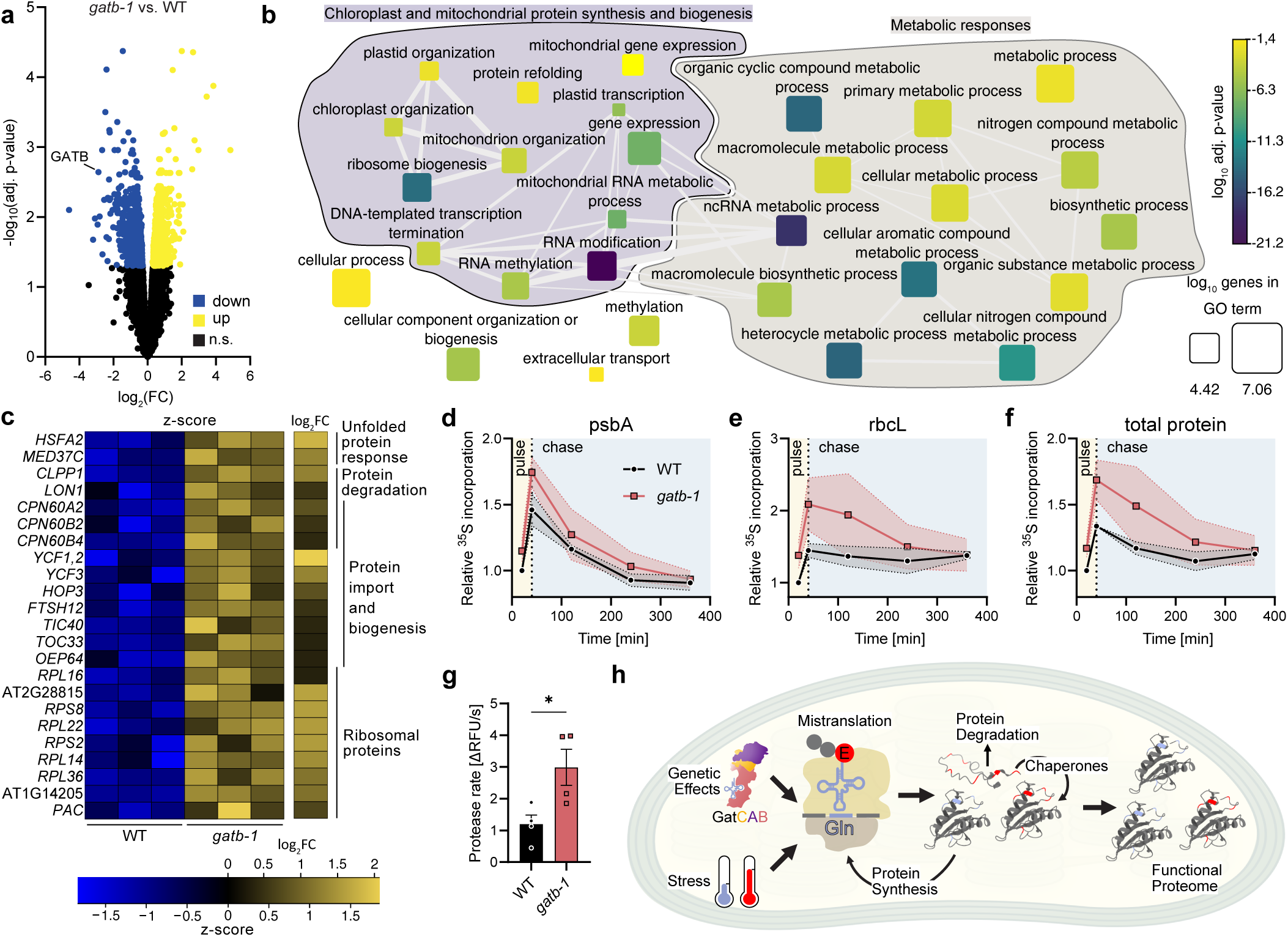
Transcriptomic profiling reveals adaptive mechanisms in response to Q→E amino-acid misincorporations in *gatb-1*. (a) Volcano plot showing differentially expressed genes (DEGs) in *gatb-1* when compared to WT mRNA. The log2 of the fold change (FC) is plotted against the negative log10 of the adjusted p-value. Significantly up-regulated genes are colored yellow and down-regulated genes a shown in blue (adjusted p-value < 0.05) while non-significantly regulated genes (n.s.) are depicted in black. (b) Condensed network of significantly enriched GO-terms in *gatb-1* vs. WT transcriptomics. The log10 of adjusted p-value of each GO-term (nodes) is indicated by the color code while the log10 of the size of the GO term is represented by different symbol sizes. The strongest 3% of pairwise similarities of the different GO terms are indicated by gray edges. GO terms were manually grouped as indicated by the background colors. (c) Heatmap showing the z-score and the log2FC of selected significantly (adj. p.value <0.05) upregulated chloroplast DEGs in *gatb-1*. For more plastid genes included in GO terms and genes of proteins located in mitochondria please refer to Extended Data Fig. 10c. (d-f) *In vivo* radiolabeling assays showing chloroplast protein synthesis and degradation of psbA (d), rbcL (e) or total protein (f). All values are relative to the first WT pulse value at 20 min. The time frame of the pulse with the ^35^S labeled amino acid is indicated by the yellow background and the chase phase with the light blue background. A representative gel can be found in Extended Data Fig. 11a. (g) Protease activity in WT or *gatb-1* stroma. A higher rate of relative fluorescence units indicates higher protease activity. For raw values, all controls and the loading control please refer to Extended Data Fig. 11b,c. Mean +/- SEM; n= 4; Unpaired t-test; *: p-value ≤ 0.05. (h) Model of molecular consequences of stress or genetic induced mistranslation in plastids.

Curated analysis of individual upregulated DEGs revealed genes involved in unfolded protein responses (UPR)/proteotoxic stress, protein degradation/stabilization/folding, protein import and biogenesis and ribosomal proteins in chloroplasts (Fig. 5c) and mitochondria (Extended Data Fig. 10c), indicating that Q→E misincorporations in endosymbiotic organelles trigger UPR along with protease and chaperone systems to avoid protein aggregation. Simultaneously, *gatb-1* increased transcripts for mitochondrial and chloroplast protein synthesis and import, presumably to stabilize the protein homeostasis and stoichiometry of multi-subunit complexes. A shift towards higher protein-transcript ratio (PTR; Extended Data Fig. 10d; see ^38^) in *gatb-1* further suggested that, besides altered transcript abundances, additional post-transcriptional mechanisms impact protein levels in mutants.

To verify this hypothesis, we examined plastid protein synthesis and degradation rates via time-resolved *in vivo* radiolabeling assays. Besides quantification of total protein turnover, high turnover rates allow the individual tracking of PSII reaction center (psbA) and rbcL. While psbA pulse and chase intensities were only by trend elevated in *gatb-1*, rbcL protein synthesis was clearly increased in *gatb-1* but rapidly returned to WT levels (Fig. 5d,e and Extended Data Fig. 11a). Similarly, total protein synthesis was higher in *gatb-1* but returned to WT levels after 320 min (Fig. 5f and Extended Data Fig. 11a) pointing to higher protease activity. A fluorescence-based protease assay using isolated stroma, confirmed significantly elevated protease activity in *gatb-1* compared to WT (Fig. 5g and Extended Data Fig. 11b,c).

Taken together, Q→E misincorporation in endosymbiotic organelles is compensated through triggered protein quality control responses including enhanced expression of chaperones, proteases and unfolded protein response genes. In chloroplasts, this is accompanied by increased protein synthesis and protease activity in the stroma.

## Discussion

Our results from Arabidopsis reveal that plants possess high tolerance towards mistranslation in mitochondria and plastids. We genetically induced significant Q→E misincorporation (≤ 91%) through decreasing GatCAB activity i.e., the indirect pathway to produce Gln-tRNA^Gln^ also common in bacteria. To the best of our knowledge, these are the highest misincorporation rates in any organism reported thus far. Overall, lack of GatCAB resulted in slower growing *gatb-1* plants, with modestly reduced photosynthetic capacity. Complete loss of GatCAB does not yield viable plants, as organellar protein synthesis cannot pursue in total absence of Gln-tRNA^Gln^. The different Q→E rates in MTex (low) and PLex (high) proteins may suggest distinct mistranslation responses in each organelle type. Since studies on translation fidelity in plastids are virtually non-existing, we focused our efforts on building an initial mechanistic model on the organelle’s response (Fig. 5h). Mistranslating plastids respond with a strong increase in protein synthesis. This yields a larger pool of stochastically mistranslated proteins. The mutants’ high protease activity and increased expression levels of chaperones allows to reach a threshold of sufficiently folded proteins to carry out their assigned functions. In contrast, bacteria and yeast mutants with error-prone mitochondrial protein synthesis have similar or lower translation rates, respectively, than wild-type cells ^39–41^. In mammalian mitochondria, even the risk of Q→E mistranslation causes a complete arrest in protein synthesis ^2,39,42^. In the future, the strategy by which plant mitochondria yield less Q→E misincorporations than plastids should be revealed. Since the transcriptomic responses for plastids and mitochondria were similar, we, for now, conclude that the mitochondrial amino-acid misincorporation compensation response in plant cells is more strict/efficient.

Chloroplasts accept highly Q→E substituted protein complexes in the photosynthetic machinery. However, the tolerance threshold varies between proteins. While Cyt*b*_6_*f* was very lenient to Q→E (≤ 91%), ATP-syn (≤ 7%) was on the other end of this spectrum. The reasons behind this bias are unclear but individual protein complex architecture may play a role. It seems surprising that *gatb-1* plants photosynthesize sufficiently while coping with mistranslation. However, several Arabidopsis mutants with defects in CO2-fixation and ATP production activate a similar cascade of protective mechanisms as we documented in *gatb-1* ^26,43–46^. *gatb-1* plants had constitutively high NPQ and accumulate xanthophylls. Together this effectively dissipates surplus energy at the PSII antenna as heat and prevents ROS formation. The main NPQ component qE and zeaxanthin formation is triggered by ΔpH across this thylakoid membrane ^47^. *gatb-1* plants have a luminal H^+^ surplus, because the low ATP-syn abundance consumes fewer H^+^. This culminates in poor ATP production, which together with 23% reduced CO2-fixation by mistranslated RuBisCO leads to decreased CO2-fixation. The resulting ATP to NADPH imbalance triggers CET and simultaneous H^+^ pumping via Cyt*b*_6_*f*. Consequentially, linear electron transport via Cyt*b*_6_*f* is decreased to protect PSI. Altogether, the protective mechanisms keep photoinhibition of PSII (decrease in *F*_v_/*F*_m_) at a minimum and enable *gatb-1* mutants to grow under ambient conditions.

Interestingly, Q→E substituted CPex proteins emerge also in wild-type plants exposed to temperature stress. Whether this is an uncontrolled response e.g., via temperature-dependent GatCAB activity changes or controlled e.g., by gene expression needs to be resolved in the future. Q→E hyper- mistranslating *gatb-1* plants responded very differently to temperature stress than WT. Wild-type plants successfully acclimated 7 days into the cold once they restored rbcL and psbA level to buffer decreased CO2-fixation and concomitant PSII photoinhibition ^48^. However, *gatb-1* plants appear unable to prevent PSII damage since newly synthesized RuBisCO complexes, with 12.4% Q→E substitutions, are malfunctioning and cannot compensate the decrease in CO2-fixation. In addition, up to 25% Q→E substitutions in PSII subunits, including psbA, may hinder PSII repair in mutants. Notably, mistranslation rates may increase during temperature stress.

Interestingly, for a 5-day-period after heat shifts, *gatb-1* performed better (higher *F*_v_/*F*_m_) than at starting ambient conditions. One explanation is that the mistranslations stress, which triggers expression of heat shock proteins and chaperones (Fig. 5c; ^11^), stabilizes protein functions and initially prevents PSII photoinhibition. A similar heat stress-tolerance mechanism was described for mistranslating bacteria ^31^. However, in plants this effect is not persistent and even the return to ambient conditions prevented an effective re-acclimation in *gatb-1*.

Our initial study of plant organellar mistranslation and genetic means to alter it stipulates new research directions. Whether naturally occurring mistranslation in plant organelles has beneficial functions during stress acclimation should be investigated. Also, evolutionary aspects should be considered. Recently, an intriguing idea was introduced: mistranslation is an important source of nongenetic variation that affects adaptive evolution on fitness landscapes ^49,50^. This hypothesis and its potential importance for plant cell evolution could be tested through experiments utilizing this and other hyper-mistranslating mutants, which await their discovery.

## Supporting information

Supplementary Table 1_Plastid Peptides wih Q-E

Supplementary Table 2_LFQ results of chloroplast proteomics dataset

Supplementary Table 3_LFQ results of amiA vs WT_amiA_dataset

Supplementary Table 4_RNAseq_results

Supplementary Table 5_Significantly enriched GO-terms

Supplementary Table 6_Sequences or all oligonucleotides

Supplementary File 3 RNAseq_Stats_and_QC

Supplementary File 1 WiscLoxHS_LB

Supplementary File 2 WiscLoxHS_RB

## Methods

### Material availability

All materials generated in this study will be made available on request, but we may require a completed materials transfer agreement.

### Growth conditions for plants

Except the plYFP lines, all *Arabidopsis* plants used in this study are the Col-0 ecotype. Seeds were surface sterilized with 70% ethanol and subsequently placed on ½ Murashige and Skoog (½ MS) basic medium ^51^ (Duchefa Biochemie) supplemented with 0.8% Phytoagar (Duchefa Biochemie), kept at 4°C in the dark for at least two days and transferred to a growth chamber (16 h at 22°C and 100 µmol photons m^-2^ s^-1^ with 30% humidity / 8 h at 18°C and 0 µmol photons m^-2^ s^-1^ with 30% humidity). After 7-9 days of growth, seedlings were transferred to soil and the grown another 14-16 days at (16 h at 22°C and 100 µmol photons m^-2^ s^-1^ with 55% humidity / 8 h at 18°C and 0 µmol photons m^-2^ s^-1^ with 65% humidity), except otherwise mentioned. For the cold and heat stress experiment, 20 days old plants were either transferred to 8°C (16 h at 8°C and 150 µmol photons m^-2^ s^-1^ with 55% humidity / 8 h at 8°C and 0 µmol photons m^-2^ s^-1^ with 65% humidity) or 37°C (16 h at 37°C and 100 µmol photons m^-2^ s^-1^ with 50% humidity / 8 h at 8°C and 0 µmol photons m^-2^ s^-1^ with 50% humidity) and grown for another 10 days after which they were transferred back to 22°C in the initial growth chamber.

The plants which were used to measure RuBisCO activity were grown as described in the following: Four to five *A. thaliana* seeds (WT, *gatb-1* and comp. line 8.3) per pot were germinated in a 0.5 L drainage pot with soil (Sunshine® mix LC-1; Sun Gro Horticulture®) in a controlled environmental chamber (Enconair Ecological Chambers Inc.) set to a 12 hours daily photoperiod, maximum PPFD of 300 µmol photons m^2^ s^-1^, and air temperatures of 23/18°C during light/dark periods. Once the seedlings germinated, all seedlings except one were removed which was grown for further analysis. The plants were watered twice a week and fertilized once a week with Sprint 330 iron chelate (1.3 g L^-1^), magnesium sulfate (0.6 g L^-1^), Scotts-Peters Professional 15-30-20 compound (2.8 g L^-1^), and Scott-Peters Soluble Trace Element Mix (8.0 mg L^-1^; Scotts).

### Mutant Plant lines

Mutant seeds for *GatA* (AT3G25660), *GatB* (At1G48520) or *GatC* (AT4G32915) were ordered from the European Arabidopsis Stock Centre (NASC). *gata-1* (SALK_08643), *gatb-2* (SALK_113005) and *gatc- 1* (GK-036F07) first generation seeds were screened for the T-DNA insertion by PCR on isolated genomic DNA (see Supplementary Table 6 for primer sequences). Insertion sites of T-DNAs were confirmed by sequencing of the PCR products. Second generation plants were screened for homozygous insertion by genotyping PCRs.

*gatb-1* (Wiscseq_DsLoxHs083_12A.1) first generation seeds were screened for the T-DNA insertion by PCR on isolated genomic DNA. Plants with homozygous T-DNA insertion in the 5’-UTR assessed by PCR showed clearly decreased photosystem II (PSII) yields (*F*_v_/*F*_m_) in Pulse-Amplitude-Modulation (PAM) Fluorometry. To exclude potential effects of secondary site mutations in the initial stock T-DNA insertion line we backcrossed a plant homozygous for the T-DNA insertion in the *GatB* 5’-UTR with a WT plant, propagated seeds of the next generation (F1) and isolated genomic DNA of second generation plants (F2) exhibiting decreased *F*_v_/*F*_m_. Isolated gDNA (NucleoSpin Plant II Mini kit, Macherey & Nagel) was subjected to 150 bp paired-end whole genome sequencing (carried out by Novogen). On the Galaxy platform ^52^, raw reads were filtered using Trimgalore with standard settings and subsequently checked using FastQC. As we were not able to find the exact border sequences of the T-DNA insertion of WiscLoxHS plant lines we initially aligned raw reads to the Tair10 genome release using Bowtie2 ^53^ with “sensitive local” settings. Aligned reads around the putative T-DNA insertion site were inspected using the IGV platform ^54^. The T-DNA insertion was confirmed to be just 5’ of the 5’-UTR of *GatB* and led to a 30 bp deletion. No reads spanned or aligned on the 30 bp deletion confirming the T-DNA insertion site and that all plants exhibiting the phenotype are homozygous for the T-DNA insertion *GatB*. By adding the left and right clipped sequences of reads aligning to the *GatB* gene to a new fasta entry of our reference genome we were able to extract an initial insertion sequence. Through iterative mapping of reads to the insertion fasta entry and adding clipped sequences to the insertion sequence were able to assess 415 bp and 549 bp of the left and the right borders of the insertion (Supplemental Files S1 and S2), respectively. We then inspected the reads which were aligned to T-DNA insertion and were clipped the 5’- and 3’-end of the T-DNA insertion. All 16 clipped reads aligned to *GatB* indicating that this T-DNA is only inserted at on locus in the genome. Additionally, we analyzed the *gatb-1* whole genome sequencing with the TDNAScan software ^55^ using the assembled insertion sequence. The results confirmed a single T-DNA insertion at 5’-UTR of *GatB* with a frequency of 1.

*gatb-1* and WT plYFP lines were created by crossing a plastid YFP expressing trans-plastomic line in the C24 ecotype ^21^ with *gatb-1*. We then isolated plants in the third generation after crossing (F3) which carry homozygous mutant or WT alleles for the *gatb-1* mutation (plYFP-*gatb-1* or plYFP-WT, respectively).

The *stn7-1* line has been established before ^28^.

### Generation of transgenic plants

For the GatB constructs used to transform Arabidopsis plants, the *GatB* coding sequence was amplified from Arabidopsis cDNA by PCR and then subsequently cloned by InFusion (Takara) into modified pGreenII vectors (based on ^56^) resulting in GatB-TEV-mVenus constructs either driven by the endogenous GatB promoter (1.8 kb upstream of the start codon of the gene) or by the strong UBQ10 promoter ^57^: pG2-UBQ10-GatB-TEV-mV_Hyg was cloned first and the UBQ10-promoter was replaced by pGATB by amplifying the promoter on gDNA and the inserting it into the vector by InFusion cloning resulting in pG2-pGATB-GatB-TEV-mV_Hyg.

To clone the amiRNA against the *GatA* subunit, amiRNA sequences were designed using the Web MicroRNA Designer, WMD3 (http://wmd3.weigelworld.org) ^58^ and subsequently cloned into a modified pGreen vector ^59^ using DNA oligos replacing the initial amiRNA and amiRNA* sequences by InFusion (Takara) cloning resulting in the vector pG2-amiRNA-GatA.

All vectors were confirmed by sanger sequencing. For plant transformation, vectors were transformed into Agrobacteria tumefaciens GV3101 ^60^ carrying the pSOUP helper plasmid by electro-poration. Successfully transformed Agrobacteria were selected on LB-Agar plates containing Rifampicin (10 µg/ml), Gentamycin (10 µg/ml) and Kanamycin (50 µg/ml). Individual colonies were checked for the presence of the respective plasmid by colony PCR using the primer which were used for the respective cloning. Positive colonies were amplified in LB medium supplemented with Rifampicin (10 µg/ml), Gentamycin (10 µg/ml) and Kanamycin (50 µg/ml). *gatb-1* or Col-0 plants were then transformed using the floral dip method ^61^. Transgenic plants were selected on 1/2 MS-Agar plates supplemented with Hygromycin (50 µg/ml). The generation and zygosity of plants used for each experiment is indicated in the text or the figure legends.

### Total leaf protein extraction

Total leaf protein was extracted by disrupting leaf material directly in precooled extraction buffer (50 mM Tris pH 7.5, 150 mM NaCl, 2.5 mM EDTA, 10% Glycerol, 0.5% NP-40, 5 mM DTT, 1x plant protease inhibitor cocktail) using a plastic pistil. Alternatively frozen and ground plant powder was resuspended in precooled extraction buffer. Cell debris was removed by centrifugation at 25,000 *g* and 4°C for 10 min. The supernatant was used for downstream applications. For equal loading protein extracts were normalized using Bradford assays.

### PAM analyses

Unless otherwise mentioned, photosynthetic parameters were measured on 3 weeks old and dark-adapted plants (15 min) plants either using the IMAGING - or DUAL-PAM (WALZ, Effeltrich, Germany). Standard induction curves were carried out at 53 µmol photons m^−2^ s^−1^. To determine the post-illumination chlorophyll fluorescence rise (PIFR) according to ^24^, plants were subjected to 56 µmol photons m^−2^ s^−1^ for 5 min, and subsequently the AL was switched off to measure the chlorophyll fluorescence Ft for additional 4 min. To measure thylakoid membrane proton conductivity (gH^+^), which is indicative ATPase activity, detached leaves were measured at 110 µmol photons m^−2^ s^−1^ and room temperature using the Photosynq MultispeQ V2.0 (PhotosynQ Inc., East Lansing, MI 48823 USA) [73] system and the RIDES 2.0 protocol.

### Immunoblot Analyses

The source of proteins samples used in each immunoblot analysis is indicated in the figure legend. Protein extracts were separated by size using SDS-PAGE, subsequently transferred on PVDF membranes (Milipore) by wet blotting. The membranes were then blocked using 5% milk powder in TBS- T (0.05% Tween-20) buffer for 1h at room temperature and then incubated with the primary antibody overnight at 4°C, subsequently washed with TBS-T, incubated with secondary antibody for 45 min at room temperature, washed again with TBS-T and visualized using a homemade ECL in an ImageQuant (GE Healthcare). Resulting tiff files were imported into Fiji ^62^, adjusted (always entire blots) and exported as .jgp files. Primary antibodies used were: α-GatAB (^3^), α-psaF (Agrisera, Prod. #: AS06 104), α-psaD (Agrisera, Prod. #: AS09 461), α-petA (Agrisera, Prod. #: AS08 306), α-petB (Agrisera, Prod. #: AS18 4169), α-psbB (Agrisera, Prod. #: AS04 038), α-psbA (Agrisera, Prod. #: AS05 084), α-psbC (Agrisera, Prod. #: AS11 1787), α-atpD (Agrisera, Prod. #: AS10 1590), α-actin (Agrisera, Prod. #: AS13 2640) and α-atpF (Agrisera, Prod. #: AS10 1604).

### Label-free quantitative t-RNA dependent trans-amidation assay

#### Expression and purification of proteins

The coding sequences without transit peptide for *GatA* (Q45-A537), *GatB* (S60-D550), *GatC* (R50-E155) and the *ND-GluRS* (At5g64050; G51-T570) from Arabidopsis were amplified by Phusion PCR (Thermo Fisher Scientific) from cDNA. The sequences for *GatA*, *GatB* and *GatC* were assembled into an artificial *GatCAB* operon detailed in ^63^ using Golden Gate Cloning (New England Biolabs). The DNA fragments encoding *GatCAB* and *ND-GluRS* were then cloned between XhoI and NcoI restriction sites in modified pET-28a vector with an improved Shine-Dalgarno sequence ^64^ resulting in genes coding for fusion proteins with a TEV-cleavable N-terminal His10-Twin_StrepII-Thioredoxin-tag: pEC-HSTrx-GatCAB and HSTrx-AtGluRS The plasmid carrying either GatCAB or GluRS were transformed into *E. coli* BL21 (DE3) cells and grown in TB medium containing 50 µg ml^-1^ Kanamycin at 37°C until the OD600 reached 0.6. Protein expression was then induced by the addition of isopropyl ß-D-1-thiogalactopyranoside (IPTG) to a final concentration of 0.5 mM and the cultures were transferred to 18 °C for 16 h. The cells were harvested by centrifugation at 3,500 *g* for 15 min at RT, resuspended in buffer A (50 mM Tris-HCl pH=8.0, 300 mM NaCl, 5% Glycerol, 2 mM DTT), flash frozen in liquid nitrogen, thawed on ice and lysed in a tissue lyser. After removing cell debris by centrifugation at 45,000 *g* and 4°C for 30 min 100 µg RNase A was added to the supernatant and incubated for 30 min on ice. All subsequent steps were carried out on ice or at 4°C with pre-cooled buffers. HIS-tagged proteins were the isolated by immobilized metal-ion affinity chromatography (IMAC) using a HisTrap HP column (GE Healthcare) on an Aekta PURE FPLC. Unbound proteins were removed by washing the column with buffer A supplemented with 20 mM imidazole and HIS-tagged proteins were eluted with 300 mM imidazole in buffer A.

The buffer was then exchanged to buffer B (50 mM Tris-HCl pH = 8.0, 150 mM NaCl, 5% Glycerol, 1 mM DTT) using a FPLC equipped with a HiPrep 26/10 Desalting column (GE Healthcare). The N-terminal tag was cleaved off by incubation with TEV protease for 16 h on ice. The TEV protease and cleaved off tag was removed by a reverse IMAC. Cut protein containing flow through was concentrated (Amicon Ultra-15, 30 kDA falcons) followed by size exclusion chromatography using a HiLoad 16/16 200 pg column (GE Healthcare) equilibrated with SEC buffer (50 mM Tris-HCl pH = 8.0, 150 mM NaCl, 10% Glycerol, 1 mM DTT). Collected fractions were analyzed by SDS-PAGE. Recombinant protein containing fractions were combined, concentrated to 2 mg/ml using Amicon Ultra-15 30 kDA falcons, aliquoted, flash frozen in liquid nitrogen and stored at -80°C.

#### Aminoacylation and tRNA-dependent trans-amidation assay

The standard reaction mixture of 50 µl was prepared containing 30 mM HEPES-KOH buffer (pH 7.0), 30 mM KCl, 15 mM MgCl2, 10 mM Gln, 10 mM Glu, 10 mM ATP, 100 µM E. coli MRE 600 total tRNA (Roche), and either 2 µM recombinant proteins or normalized (by Bradford assays; Bio-Rad) Arabidopsis stromal extract similar to ^3^. This mixture was then incubated for the indicated time at 25°C. Reactions were stopped by addition of sodium acetate (pH 5.0) to a final concentration of 1.5 M. Subsequently, RNA was cleaned-up using the Monarch RNA Cleanup Kit (New England Biolabs) and eluted in 100 µl of RNase-free water, quantified with a Nanodrop, and concentrations were adjusted to 1.25 µg/µl (∼50 µM) tRNA. De-acylation was carried out by adding KOH to a final concentration of 30 mM and incubation for 30 minutes at 37°C, followed by neutralization with 30 mM HCl. To measure glutamate and glutamine by luminescence, samples were prepared in a 96-well plate according to the manufacturer’s instructions (Glutamine-Glutamate-Glo Assay; Promega). The resulting luminescence was measured using a Spark Multimode Microplate Reader (Tecan) with an integration time of 1,000 ms per count.

### Protoplast isolation and confocal microscopy for protein localization studies

The protoplast isolation was conducted according to the “Tape-Arabidopsis-Sandwich” method ^65^. Leaves measuring approximately 3 cm in length and 1 cm in width were harvested from 4-week-old plants grown under optimal light conditions (∼ 120 µmol photons m^-2^ s^-1^). The upper epidermal surface of each leaf was secured to a strip of time tape (Tesa) for stabilization, while the lower epidermal surface was attached to a strip of scotch tape [Tesa]. The Scotch tape was then carefully removed, stripping away the lower epidermal layer while the leaf remained adhered to the Time tape. The peeled leaves (4 to 5 leaves) still attached to the Time tape were placed in a petri dish containing 10 ml of enzyme solution (1% cellulase “Onozuka R-10” (SERVA), 0.25% Macerozyme R-10 (SERVA), 0.4 M mannitol, 10 mM CaCl2, 20 mM KCl, 0.1% BSA and 20 mM MES, pH 5.7). The petri dishes were gently shaken on a platform shaker at 20 rpm in darkness for 60 min until the protoplasts were released into the solution. The protoplasts were then collected by centrifugation at 100 *g* for 3 min and washed twice with pre-chilled W5 modified wash solution (154 mM NaCl, 125 mM CaCl2, 5 mM KCl, 5 mM Glucose, and 2 mM MES, pH 5.7). Finally, the protoplasts were centrifuged again and resuspended in 200 µl of modified MMg solution (0.4 M Mannitol, 15 mM MgCl2, and 4 mM MES, pH 5.7).

Images were acquired using a Leica Stellaris 5 Confocal Laser Scanning Microscope, featuring a supercontinuum laser (Wight Light Laser & 405 nm diode) and a silicon-based Multi-Pixel Photon Counter (MPPC) hybrid detector. For protein localisation studies, yellow fluorescent protein (YFP) was excited at a wavelength of 514 nm while chlorophyll autofluorescence (chl *a*) was excited at 405 nm. Emission was detected at 520–580 nm for YFP and 623–813 nm for chlorophyll. A Z-stack of 11 images was acquired and processed using the Las X software to create the maximum projection.

### HPLC based pigment quantification

Pigments were extracted from frozen leaf material (30-50 mg) upon addition of 100% ice-cold acetone using a Heidolph RZR 2051 homogenizer (1,000 rpm, for about 1 min). Samples were centrifuged for 5 min at 17,000 *g* (4°C) and the resulting supernatant was filtered (pore size 0.2 μm). Pigments were separated and quantified by high-performance liquid chromatography (HPLC) as described (Färber et al. 1997).

### Transmission-electron microscopy

After cutting WT and *gatb-1* leaves into 1 mm^2^ pieces in fixation buffer (75 mM cacodylate; 2 mM MgCl2 and 2.5% glutaraldehyde) the buffer was infiltrated into the leaf pieces by alternating the surrounding pressure. Leaf samples were then incubated overnight at 4°C and subsequently post-fixed in 1% osmium tetroxide for 1 h. After, samples were dehydrated by consecutively incubating the samples in 10% acetone for 15 min, 20% acetone supplemented with 1% uranyl acetate for 30 min, and in 40, 60, and 80% acetone for 20 min each concentration, 100% acetone for 5 min and the nonce more overnight in 100% acteone. Then, Spurr’s resin was infiltrated and polymerized at 63°C and subsequently thin sectioning was performed. Transmission-electron microscopy was carried out using a JEOL F200 with 200 kV (JEOL, Freising, Germany). Images acquisition was done using a bottom-mounted Xarosa 20 mega-pixel CMOS camera (EMSIS, Münster, Germany).

### Chloroplast enrichment

21-24 days old plants used for the chloroplast enrichment were grown at the same time at standard growth conditions defined in the “experimental model and study participant details” section. To reduce starch accumulation in the chloroplasts, plants were kept dark for 16h before chloroplast enrichment. Chloroplasts were enriched by disrupting ∼3 g leaves in cold CP-MT extraction buffer (0.3 M Sorbitol, 5mM MgCl2, 5 mM EDTA, 20 mM HEPES-KOH pH 8.0 and 10 mM NaH2CO3) using a polytron mixer (Bachofer Laboratoriumsgeräte). The resulting suspension was filtered through two layers of Miracloth. Chloroplasts were then pelleted by centrifugation at 1,500 *g* and 4°C for 5 min. The chloroplast containing pellet was then washed with chloroplast extraction buffer, pelleted again and then used in subsequent analyses.

### Chloroplast fractionation and stroma preparation

To lyse enriched chloroplasts (see “Chloroplast enrichment”) the chloroplasts were resuspended in chloroplast burst buffer (10 mM Hepes pH 7.5, 10 mM MgCl2) and passed through a syringe needle 15- 20 times. Subsequently, NaCl was added to gain a final concentration of 150 mM. To remove thylakoid membranes the suspension was centrifuged at 5,000 *g* and 4°C for 5 min. Supernatant was transferred to a fresh tube and residual liquid was carefully removed. The thylakoid containing pellet was resuspended in 8 M Urea and flash frozen in liquid nitrogen (thylakoid fraction). To further remove residual thylakoid, inner and outer envelope membranes as well as aggregates, the supernatant was subsequently centrifuged twice at 45,000 *g* and 4°C for 15 min. The supernatant (stroma fraction) was either directly used for downstream analyses or flash frozen in liquid nitrogen in aliquots.

### RuBisCO activity assay and quantification

Leaf punches (surface area ∼ 2.2 cm^2^) were collected from plants that are at least six weeks old and immediately frozen in liquid nitrogen prior to analysis. Frozen leaf tissue was ground using mortar and pastel with 0.5 ml of extraction buffer [100 mM EPPS-NaOH [pH 8.0]; 1% [w/v] polyvinylpolypyrrolidone [PVPP]; 1 mM ethylenediaminetetraacetic acid (EDTA) [pH 8.0]; 10 mM dithiothreitol [DTT]; 0.1% [v/v] Triton X-100; 5 mM MgCl2; 20 μL plant protease inhibitor (PI) (Sigma)]. Ground samples were centrifuged at 13,300 *g* for one minute and RuBisCO enzyme was activated with 50 mM NaHCO3 and 40 mM MgCl2 for at least 45 minutes at room temperature.

RuBisCO activity was determined by adding 20 µl of activated enzyme extract in a 0.6 mL assay buffer [100 mM EPPS-NaOH (pH 8.0); 20 mM MgCl2; 1 mM EDTA; 0.5 mM DTT; 1 mM adenosine triphosphate (ATP), 5 mM phosphocreatine; 0.2 mM NADH; 20 mM NaHCO3, 25 units ml^-1^ creatine phosphokinase, 250 units mL^-1^ carbonic anhydrase, 25 units ml^-1^ 3-phosphoglyceric phosphokinase, 25 units ml^-1^ glyceraldehyde 3-phosphate dehydrogenase, 200 units mL^-1^ triose-phosphate isomerase, 20 units ml^-1^ α-glycerophosphate dehydrogenase]. The reaction was initiated by adding 0.6 mM RuBP and measured by monitoring the consumption of NADH at 340 nm wavelength using a spectrophotometer (Evolution 300 UV-VIS, Thermo Fisher Scientific) at 25°C and 35°C.

RuBisCO active sites were quantified from the stoichiometric binding of radiolabeled [^14^C]carboxyarabinitolbisphosphate (^14^CABP) ^66^. An aliquot (100 μl) of activated plant extract was incubated with 5 µl of 2 mM ^14^CABP for 45 min before it was gravity eluted on a chromatography column (7374731; Bio-Rad) packed with Sephadex G-50 fine beads (GE Healthcare Biosciences). The fractions of RuBisCO-bound ^14^ CABP were collected using the elution buffer [20 mM EPPS-NaOH [pH 8.0], 75 mM NaCl] and analyzed using a liquid scintillation counter (Tri-Carb 2100TR, Perkins-Elmer).

### Blue-Native PAGE

BN-PAGE was carried out according to Nickel et al, 2016 ^67^. Enriched chloroplasts (see above) were used immediately after extraction. Here, the chlorophyll concentration of all samples was assessed in 80% acetone using a photometer. Subsequently, the volume corresponding to 30 µg chlorophyll was used for all samples for the BN-PAGE used for mass spectrometry. *gatb-1* only has 73% of the Chl *a* + *b* content (in μmol/g FW) of WT plants (see Fig. 1c). Thus, to normalize the loading for the analytical BN-PAGE to the fresh weight, the volumes for 30 µg for WT and 21.9 µg *gatb-1* were used. Chloroplasts for both approaches were then pelleted at 500 *g* and 4°C for 3 min, resuspended in ACA buffer (50 mM Bis-Tris pH 7.5, 750 mM Aminocaproic acid and 0.5 mM EDTA-Na2), supplemented with dodecyl maltoside to a final concentration of 0.5% (w/v), incubated on ice for 25 min and finally centrifuged at 16,000 *g* and 4°C. BN-loading dye (final concentrations: 68 mM Aminocaproic acid and 0.45% (w/v) Coomassie brilliant blue G-250) was added to the supernatant before loading half of the samples on 4- 12% Acrylamide gels (final concentration of gel buffer: 500 mM Aminocaproic acid and 50 mM Bis-Tris pH 7.0). The gel was run using blue cathode buffer (50 mM Tricine, 15 mM Bis-Tris and 0.2% Coomassie brilliant blue G-250) and anode buffer (50 mM Bis-Tris pH 7.0) until the sample dye front reached 2/3^rd^ of the gel. Then the cathode buffer was changes to clear cathode buffer without Coomassie brilliant blue. When the dye front reached the bottom of the gel, the gel was removed from the glass plate and imaged. A part of the gel was then cut and stained with Coomassie brilliant blue R-250.

### Mass spectrometry

#### Protein extraction and sample preparation amiGatA leaf MS dataset

As *amiGatA* plants are very small, extraction of chloroplasts from these plants was not feasible. Thus, proteins were extracted from above ground rosettes. To that end, 145 T2 plants of *amiGatA* were grown alongside WT and *gatb-1*. After 28 days of growth, per biological WT and *gatb-1* replicate 4 plants as well as per *amiGatA* biological replicate 35 rosettes were pooled and immediately flash frozen in liquid nitrogen. A total of 4 biological replicates per genotype was prepared. Frozen plant material was ground to fine powder using a precooled mortar and pestle. 100 mg of the powder was resuspended in 0.3 ml of extraction buffer (100 mM HEPES pH 7.5, 150 mM NaCl, 10 mM DTT, 6 M Guanidine Chloride, Roche cOmplete™ Protease Inhibitor Cocktail). Each sample underwent sonication using a Branson Sonifier B-12 (Branson Ultrasonics, Danbury, USA) for 3 cycles of 20 sec and samples were subsequently incubated at 60°C for 10 min. Cell debris was removed by centrifugation at 10,000 *g* for 15 min. Proteins were precipitated with chloroform-methanol. For this, methanol/chloroform and water were added to the clarified sample in a 4:2:3 ratio. The mixture was centrifuged for 10 min at 10,000 *g*, the precipitated protein accumulating at the interphase was retained and washed five times with methanol. The protein pellet was dried and solubilized in 3 M Urea/2 M Thiourea in 50 mM HEPES (pH 7.8). The total protein concentration was determined using the Pierce 660 nm Protein Assay Kit (Pierce, Thermo Fischer Scientific). 100 µg of protein was reduced with 10 mM DTT for 30 min at 37°C and alkylated with 50 mM iodoacetamide (IAA) for 20 min at RT in darkness, followed by overnight digestion with 1 µg Trypsin (Pierce Thermo Fischer Scientific), at 37°C.

#### Protein extraction, sample preparation L_8_S_8_ *RuBisCO* and chloroplast MS datasets

Plants (WT, *gatb-1* and comp line 8.3) used for the chloroplast MS dataset were grown at the same time and the chloroplast enrichment (see “Chloroplast enrichment”) was carried out in parallel on one day. For the L_8_S_8_ RuBisCO dataset, the chloroplast enrichment and stroma isolation (see “Chloroplast fractionation and stroma preparation”) with subsequent size exclusion chromatography (SEC) to isolate L_8_S_8_ RuBisCO for all bio replicates was done consecutively with alternating genotypes within two days. A total of 4 biological replicates per genotype was prepared.

To isolate fully assembled L_8_S_8_ RuBisCO complexes, isolated stroma was immediately subjected to size exclusion chromatography (SEC). Here, a Superdex 200 Increase 10/300 GL column was equilibrated with SEC buffer (10 mM Hepes pH 7.5, 150 mM NaCl, 4% Glycerol) and then 300 µl of isolated stroma were injected and eluted. 500 µl fractions were collected. Fractions 9-12 of each run were combined and concentrated to 70-100 µl using a centrifugal concentrator (MWCO 100 kDa; Vivaspin).

Enriched chloroplasts or concentrated SEC fractions (equivalent to 50 µg protein) were resuspended in 250 µl extraction buffer (6 M Guanidine Chloride, 0.1 M HEPES pH 7.6), sonicated, incubated at 60°C and centrifuged as described above for plant powder. Trypsin digestion was performed by the FASP method according to ^68^. In brief, the clarified supernatant was added to a Microcon-30kDa centrifugal filter Unit (Merck Millipore, Burlington, USA) and centrifuged at 14,000*g* for 20 min. The filter units were washed with 8 M urea in 0.1 M HEPES (pH 7.8) and proteins were reduced with 10 mM DTT for 10 min at 37°C. After centrifugation proteins were alkylated with 0.05 M IAA for 5 min in darkness, again followed by centrifugation. Filter units were washed with 8 M urea in 0.1 M HEPES (pH 7.8) and subsequently with 0.1 M HEPES (pH 7.8). Digest was performed with 0.5 µg Trypsin (Pierce Thermo Fischer Scientific) at 37°C overnight. To elute the digested peptides, filter units were centrifuged, 50 µl of 0.5 M NaCl were added and again centrifuged.

#### Sample preparation BN-PAGE and plYFP MS datasets

Excised Coomassie-stained gel pieces from BN- or SDS-PAGE gels were washed three times with water and cut into small pieces. Gel pieces were washed twice for 10 min each with 20 mM ammonium bicarbonate (ABC) and acetonitrile (ACN). Subsequently, proteins were reduced with 10 mM DTT at 56°C for 30 min. Samples were washed with ACN for 10 min followed by alkylation with of 55 mM IAA for 30 min in darkness. The samples were washed twice for 10 min each with ABC and ACN, and then digested overnight with 0.3 μg Trypsin (Pierce Thermo Fischer Scientific) at 37°C. Digested peptides were extracted with 5% formic acid (FA), 50% ACN, 50% ACN and 1% FA, each for 15 min at 37°C. All extractions were pooled and peptides were dried in a vacuum centrifuge.

#### LC-MS/MS analyses of chloroplast, amiGatA leaf, BN-PAGE, plYFP, and L_8_S_8_ *RuBisCO* MS datasets

Digested peptides from all above protocols were purified using home-made C18 stage tips ^69^. The tips were activated with 100% methanol and equilibrated with 0.5% FA before loading the peptides. After loading, the tips were washed with 0.5% FA. Peptides were eluted with 80% ACN with 0.1%FA.

For LC-MS/MS analysis 1 µg of peptides were separated over linear gradients from 5% to 80% (v/v) ACN at a constant flow rate of 250 nl/min on a nano-LC system (Ultimate 3000 RSLC, Thermo Fisher Scientific, Waltham, MA, USA). Gradient lengths varied according to sample complexity: total protein sample 90 min, chloroplast sample 60 min, gel sample 30 min. The LC was equipped with an Acclaim Pepmap nano-trap column (C18, 100 Å, 100 μm × 2 cm) and an Acclaim Pepmap RSLC analytical column (C18, 100 Å, 75 μm × 50 cm), both from ThermoFisher Scientific. The column temperature was maintained at 50°C throughout the run. MS/MS was performed on an Impact II high-resolution Q-TOF (Bruker Daltonics, Bremen, Germany) using a CaptiveSpray nano electrospray ionization (ESI) source (Bruker Daltonics). MS1 spectra with a mass range from m/z 200–2000 were acquired at 3 Hz, and the 18 most intense peaks were selected for MS/MS analysis with an intensity-dependent spectrum acquisition rate of 4 to 16 Hz. The dynamic exclusion duration was set to 0.5 minutes.

#### Protein extraction and MS sample preparation for mitochondria MS dataset

Mitochondria from WT and *gatb-1* plants were enriched by blending ∼3 g leaf tissue of 24 day old plants in cold CP-MT extraction buffer (0.3 M Sorbitol, 5 mM MgCl2, 5 mM EDTA, 20 mM HEPES-KOH pH 8.0 and 10 mM NaHCO3) using a polytron mixer (Bachofer Laboratoriumsgeräte). The resulting suspension was filtered through two layers of Miracloth. Chloroplasts and thylakoid membranes were removed by centrifuging the filtered suspension twice at 2,500 *g* and 4°C for 5 min. Mitochondria were then enriched by centrifuging the supernatant of the previous steps at 20,000 *g* and 4°C for 20 min. The resulting pellet was washed one more time by resuspending it in CP-MT extraction buffer and centrifuging at 20,000 *g* and 4°C for 20 min. The supernatant was carefully removed the pellets were resuspended with 100 µl 4% SDS in 50 mM Tris, pH 7.5. Samples were prepared for MS analysis following the SP3 protocol ^70^, using TCEP (5 mM) and CAA (14 mM) for reduction and alkylation. Proteins were bound to beads by the addition of 100 µl EtOH and washed twice with 80% EtOH. They were digested in 50 mM TEAB, pH 8.5 at 37°C. After the overnight digestion, peptides were eluted in fractions with 30 μl of each elution buffer (89%, 86%, 82%, 0% of isopropanol) and pipetting up and down according to ^71^. Peptides were dried by vacuum centrifugation and reconstituted in 0.5% TFA and 2% acetonitrile for LC-MS/MS analysis.

#### LC-MS/MS analyses of mitochondria MS dataset

LC-MS/MS analysis was performed by using an EASY-nLC 1200 coupled to an Exploris 480 mass spectrometer (Thermo Fisher). Separation of peptides was performed on 20 cm frit-less silica emitters (CoAnn Technologies, 0.75 µm inner diameter), packed in-house with reversed-phase ReproSil-Pur C18 AQ 1.9 µm resin (Dr. Maisch). The column was constantly kept at 50°C. Peptides were eluted in 115 min applying a segmented linear gradient of 0% to 98% solvent B (solvent A 0% ACN, 0.1% FA; solvent B 80% ACN, 0.1% FA) at a flowrate of 300 nl/min. Mass spectra were acquired in data dependent acquisition mode. MS^1^ scans were acquired at an Orbitrap Resolution of 120 000 with a Scan Range (m/z) of 280-1500, a maximum injection time of 100 ms and a Normalized AGC Target of 300%. For fragmentation only precursors with charge states 2-6 were considered. Up to 20 Dependent Scans were taken. For dynamic exclusion the exclusion duration was set to 40 sec and a mass tolerance of +/- 10 ppm. The Isolation Window was set to 1.6 m/z with no offset. A normalized collision energy of 30 was used. MS^2^ scans were taken at an Orbitrap Resolution of 15 000, with a fixed First Mass (m/z) = 100. Maximum injection time was 150 ms and the normalized AGC Target 5%.

#### Cold- and heat-treated WT plants MS dataset

Raw data for the cold- and heat-treated WT plants dataset ^34^ was downloaded from the ProteomeXchange Consortium via the PRIDE partner repository ^72^ with the identifier PXD006797 (reviewer account details: Username, Password: OopPQSdq).

#### Database search for all MS datasets

Chloroplast, BN-PAGE and L_8_S_8_ RuBisCO raw data files were analyzed using MaxQuant software (v2.1.4.0) ^73^, with peak lists compared against the Arabidopsis reference proteome from UniProt (www.uniprot.org). Immunoprecipitated plYFP data was analyzed using MaxQuant software (v2.2.0.0) matching spectra to the Arabidopsis reference proteome from UniProt (www.uniprot.org) to which the sequence of the plastid expressed YFP was added. Additionally, GFP and YFP was removed from the common lab contaminants list for the search. For the mitochondria and heat/cold datasets, peak lists were compared with the Araport 11 proteome (Araport11_genes.201606-pep) to which fasta entries from an updated plastid and mitochondria proteome ^74^ were added. LFQ quantification and ’match-between-runs’ were enabled. Oxidation (M), Acetyl (Protein N-term), Asn -> Asp, and Gln -> Glu were included as variable modifications, while Carbamidomethyl (C) was set as a fixed modification. IBAQ protein quantification was enabled. All other settings were kept at default values.

#### Calculation of percentage of unique modified peptides and rate of peptides with amino-acid misincorporation from MS Data

The MaxQuant evidence output file was used to assess the number of unique modified peptides per genotype and misincorporation rate. Uniprot identifier were converted to Tair gene identifier using Perseus (v2.0.7.0) ^75^. Only peptides identified by MULTI-MS/MS were considered.

Deamidation of Q and N to E and D, respectively, is a spontaneously occurring process and would lead to the same mass shift as a Q→E mistranslation. For intact proteins deamidation rates are slow (half-life 1−500 days for N, and 100−5000 days for Q; ^76^) while deamidation rates can be much higher during the sample preparation for mass spectrometric measurements ^77^. The half-life of most chloroplast proteins is considerably shorter ^78^ than the half-life of deamidation of N and especially Q in intact proteins. Thus, *in vivo* deamidation is unlikely to accumulate in chloroplast proteins because of the high turnover. Additionally, as N is getting deamidated faster in intact proteins one would expect more deamidation on this residue when compared to Q but N→D rates are similar in mutants and WT proteins. More rapid deamidation during the sample preparation would occur for all samples in similar rates. Based on this information we consider changed Q→E in our mutants when compared to WT control plants not to be caused by differing *in vivo* deamidation rates but rather by genetic effects.

When calculating the percentage of unique peptides with amino acid misincorporation per genotype no peptide was counted multiple times in one biological replicate in case it was identified in multiple runs, fractions or with different charges. Only one identification per biological replicate was considered. Only peptides with one kind of amino-acid substitution were taken into consideration. To calculate the percentage of modified peptides carrying an amino-acid change of one kind stemming either from cytosolic, mitochondrial or plastid expression, the number of peptides carrying one kind of amino-acid misincorporation from either origin of expression was divided by the total number of unique peptides (modified or non-modified) from the same origin of expression per biological replicate.

To calculate the percentage of peptides with amino acid misincorporations, the intensities of all identifications (runs, fractions or charges) of one modified peptide per biological replicate were summed up (Int_MOD_). Then, the intensities of all different versions (modified or non-modified) of that same peptide were also summed up within the same biological replicate (Int_ALL_). Lastly, the percentage of each modified peptide was calculated by dividing the Int_MOD_ by Int_ALL_ per biological replicate similar to ^8–10^.

### Label-free quantification and bioinformatic analysis of protein abundances

Protein quantification was performed using the label-free quantification (LFQ) algorithm ^79^ within the MaxQuant Software ^73^. Further analysis was done using Perseus (v2.0.7.0) ^75^. To improve the dataset quality, potential contaminants, proteins identified only through site modification, and reverse hits were removed. Only protein groups quantifiable by the LFQ algorithm in at least three out of four replicates in at least one condition were retained. LFQ intensities were then log2transformed, and missing data were imputed from a normal distribution within Perseus using standard parameters. Differentially expressed proteins were determined in the Perseus software with standard settings. Statistical test and cutoffs are indicated in the figure legends.

### RNAseq sample preparation and sequencing

21-day old plants were harvested after 4 hours of light. To that end, whole above ground material of plants was taken and immediately flash frozen in liquid nitrogen. For each biological replicate 4 plants were combined and for each genotype plants for 3 biological replicates were harvested. Frozen plant material was homogenized into fine powder with pre-cooled mortar and pestle. Subsequently, total RNA was extracted using the RNeasy Plant Mini Kit (Qiagen, Hilden, Germany) with on column DNAse digestion according to manufacturer’s instructions. After an initial check of the RNAs using a NanoDrop (Thermo Fisher) all RNA extractions were tested on an Bioanalyzer RNA 6000 Nano assay (Agilent, Santa Clara, CA, USA). Library preparation and Illumina paired-end 150 bp sequencing was carried out by BMKGene (Münster, Germany).

### RNAseq data analysis

Data analysis of the RNAseq experiment has been carried out with standard settings on the Galaxy platform ^52^ on usegalaxy.eu, unless otherwise mentioned. Raw reads were trimmed using TrimGalore (v0.6.7), then checked using FastQC (v0.73).

For the differentially expressed genes analysis, reads were mapped on the reference genome Arabidopsis Tair 10 (Arabidopsis_thaliana.TAIR10.dna.toplevel.fa.gz downloaded from ensembl.org) using the RNAStar mapper (v2.7.8a) ^80^. After mapping quality control using the RSeQC package (v5.0.1) ^81^ read pairs (fragments) per gene were counted using featureCounts (v2.0.1) ^82^ together with TAIR 10 gene annotations (Arabidopsis_thaliana.TAIR10.52.gtf downloaded from ensembl.org). All log files and outputs of the quality control were combined into one html file (Suppl. file 3) using MultiQC (v1.11) ^83^.

To quantify reads as transcript per million (TPM) filtered reads were used as input for Kallisto (v0.48.0) ^84^ together with a Araport 11 transcriptome (Araport11_cdna_20220914_representative_gene_model.fasta).

Statistical analysis and assessment of differentially expressed genes using the read count files (see “RNAseq data analysis”) was performed with limma-voom (v3.5.0.1) including EdgeR ^85–87^. The limma-voom package was set to low read count filtering to at least 1 counts per million in at least 3 samples, TMM read count normalization, and one factor analysis. Statistical cutoffs are indicated in the figure legends.

### GO-Term enrichment and condensation

To assess enriched gene ontology-terms the gene identifier of significantly upregulated genes (adjusted p-value <0.05; see Supplementary Table 4) in *gatb-1* when compared to WT were pasted into the web portal of g:Profiler (https://biit.cs.ut.ee/gprofiler;^88^) using the *Arabidopsis* organism with default values. To condense significantly enriched GO-terms (see Supplementary Table 5) and their p-values were then used on the Revigo web server (http://revigo.irb.hr/; ^89^). The resulting network was exported, imported into cytoscape (v3.9.1) ^90^ and node were resized and recoloured according to the Revigo p-values and sizes of GO-terms.

### Calculation of correlation of protein/transcript and protein transcript ratios (PTR)

The calculation of the correlation of peptide vs. transcript and protein transcript ratios (PTR) was carried out similar to ^38^. Quantitative data for transcripts (transcript per million; TPM; see “RNAseq data analysis“) and proteins (intensity Based Absolute Quantitation ;iBAQ; see “Database search for all MS datasets”) were used. Only proteins which had valid iBAQ values for all 4 biological replicates in both, WT and *gatb-1* datasets (Chloroplast MS dataset) and genes with a median TPM >1 were considered. 1,072 transcripts/proteins met these criteria. Correlation and linear regression analysis were performed using the GraphPad Prism software.

PTR were calculated by dividing the median iBAQ values with the median TPM values per transcript/protein followed by a log2 transformation of the values. The distribution of PTRs in WT and *gatb-1* plants was calculated using the GraphPad Prism software.

### *In vivo* radiolabeling (pulse-chase assay)

10 mg leaf material from 3-week-old Arabidopsis plants grown on soil were harvested and transferred into 1.5 ml tubes with 50 µl reaction buffer (1 mM KH2PO4 pH 6.3, 75 µM cycloheximide to inhibit cytosolic translation). Samples were incubated for 30 min on ice in darkness. 30 µCi 35S- methionine/cysteine was added to each sample and the samples were infiltrated in a vacuum centrifuge for 1 min. Samples were incubated at constant temperature of 25°C and high light illumination of 1,000 µE m^-2^ s^-1^ for the indicated time points (pulse). After the incubation the reaction buffer was removed, and the samples were washed with 200 mM Na2CO3 to remove residual 35S-methionine/cysteine. For the chase, samples were incubated in reaction buffer with 10 mM unlabeled methionine and cysteine under the same conditions for indicated time points. Leaves were homogenized in 100 mM Na2CO3 and loaded on 12% SDS gels. Gels were Coomassie stained, dried and exposed to a phosphor imaging plate overnight.

### Protease activity assay

Frozen aliquots of isolated stroma (see “Chloroplast fractionation and stroma preparation”) were thawn on ice. The protease activity of 15 µl Stroma (except the “no stroma” samples) were measured in 200 µl reactions in 96 well plates according to manufacturer’s instructions (EnzChek Protease Assay Kit; Invitrogen). For the “cooked stroma” samples stroma was heated at 95°C for 10 min and centrifuged at 20,000 *g* for 5 min. The supernatant was used in the protease activity assay Fluorescence was measured in two technical replicates per biological replicate in a Spark Multimode Microplate Reader (Tecan).

### Immunoprecipitation and characterization of plYFP fluorescence

Total proteins were extracted from 500 ul of frozen ground powder of 4 biological replicates of either plYFP-WT or plYFP-*gatb-1* leaf material with 2 ml of extraction buffer (see: “total leaf protein extraction”) and kept on ice. An aliquot was taken and subjected to western blot analysis (see “Western Blot Analyses”). Protein amounts were normalized using Bradford and the YFP fluorescence intensity in was assessed in 96 well plates in a Spark Multimode Microplate Reader (Tecan).

To prepare the samples for the MS analysis of plYFP another immunoprecipitation was performed Total proteins were extracted from 500 ul of frozen ground powder of 4 biological replicates of either plYFP- WT or plYFP-*gatb-1* leaf material with 2 ml of extraction buffer (see: “total leaf protein extraction”) and kept on ice. homemade GFP-trap based on a double ankyrin-repeat protein ^91^ was added to the protein extract and the mixture was incubated on a rotation shaker for 2 h at 4°C. The resin was pelted by centrifugation at 500 *g* for 1 min and washed 4 times with wash buffer (50 mM Tris pH 7.5, 150 mM NaCl, 2.5 mM EDTA, 10% Glycerol, 0.5% NP-40). After pelleting the resin after the last wash step, the resin was resuspended in 1.5 x SDS-Loading dye, cooked at 80°C, centrifuged at 500 *g* for 1min and the supernatant was subjected to SDS-PAGE in a 4-20% Gel (Mini-PROTEAN TGX; Biorad). Gels were stained with coomassie brilliant blue and bands corresponding to YFP were cut out and subjected to MS analysis (see “Sample preparation BN-PAGE and plYFP MS datasets”)

### Visualization of protein structures

ChimeraX (v1.7.1) ^92^ was used to prepare all images from protein structures. PDB IDs and more information of the structural representation can be found in the figure legends.

### Hierarchical clustering

Hierarchical clustering, cluster profiles and resulting heatmaps were produced with Perseus (v2.0.7.0)^75^. Individual parameters are mentioned in the figure legends.

### Statistical analyses

All replicates, except the replicates using recombinant GatCAB and GluRS as the positive control in the quantitative tRNA dependent trans-amidation assay, used in this study were biological replicates. For the replicates of the aforementioned positive control of the tRNA dependent trans-amidation assay the same purifications of GatCAB and GluRS were used. All other statistical tests which were not explained above were done using in the GraphPad Prism software. The tests carried out, number of replicates and statistical cutoffs are mentioned in the figure legends for each experiment.

## Acknowledgments

We thank Prof. Dr. Ralph Bock (Max Planck Institute of Molecular Plant Physiology) for providing germplasm of the trans-plastomic YFP (plYFP) reporter line pJF1151. We are grateful for insightful discussions with Prof. Sebastian Leidel (Cellular RNA Biochemistry, University Bern). Prof. Dr. Anne-Marie Duchêne (Institut de Biologie Moléculaire des Plantes, CNRS Strasbourg) kindly provided the GatAB antibody. We thank Dr. Jörg Meurer (LMU) for valuable discussion of Imaging and DUAL PAM data and Paulina Heinkow for technical assistance, especially in maintaining the LC-MS/MS instruments at the MSPUB (WWU). We are grateful for technical undergraduate assistance by LMU alumni: Felix Pförtner and Daniel Rehm. The initial isolation of the *gatb-1* allele was supported by Washington State University alumni: Chase L. Lewis, Chance M. Lewis, and Laura S. Lopez.

H-H.K., B.B., Seb.S., and M.K. were funded by the Deutsche Forschungsgemeinschaft (DFG) (SFB-TR 175, project B09). Confocal microscopy work was funded by DFG (INST 86/2231-1 FUGG) to H-H.K.. Early project funding came from an National Science Foundation (NSF) Career Award (IOS-1553506) to H-H.K.. Ser.S. was funded by DFG SFB-TR 175 (project B06). J.E. and I.F. MS instruments were funded by DFG (INST 211/744-1 FUGG). Further support was provided by DFG FOR 5573 (project 06 H-H.K., project 09 I.F.). TEM instrument was funded by the DFG (INST 86-1852-1) to A.K.. A.B.C. was supported by the Division of Chemical Sciences, Geosciences, and Biosciences, Office of Basic Energy Sciences, Department of Energy (grant no. DE-SC0001685).

## Author Contributions

H-H.K. designed research, designed figures, and wrote the manuscript. B.B. designed research, performed most experiments, analyzed data, designed and prepared figures, and wrote the manuscript. Seb.S. established the trans-amidation assay, measured GatCAB activities, performed confocal laser microscopy and designed a figure. Ser.S. performed proteomics and pulse-chase experiments. M.K. performed most PAM measurements. C.E. established transgenic Arabidopsis plants. A.K. collected electron micrographs. P.J. analyzed leaf pigments, K.O. and A.B.C. performed RuBisCO assays. J.E. and I.F. performed mitochondria proteomics. All authors assisted in editing the manuscript.

## Supplemental Information

Supplementary Table 1: Excel file with Plastid Peptides containing Q and Q→E misincorporation rates when applicable

Supplementary Table 2: Excel file with LFQ results of chloroplast proteomics dataset

Supplementary Table 3: Excel file with LFQ results of *amiGatA* vs WT dataset

Supplementary Table 4: Excel file with RNAseq results

Supplementary Table 5: Excel file with significantly enriched GO-terms of upregulated genes in *gatb-1* vs WT transcript abundance

Supplementary Table 6: Excel file with sequences or all oligonucleotides used in this paper

Supplemental file 1: WiscLoxHS left border sequence as fasta

Supplemental file 2: WiscLoxHS right border sequence as fasta

Supplemental file 3: RNAseq quality control and stats as html file

## Corresponding authors

Correspondence should be addressed to Hans-Henning Kunz and Benjamin Brandt

---

**Extended Data Fig. 1:**
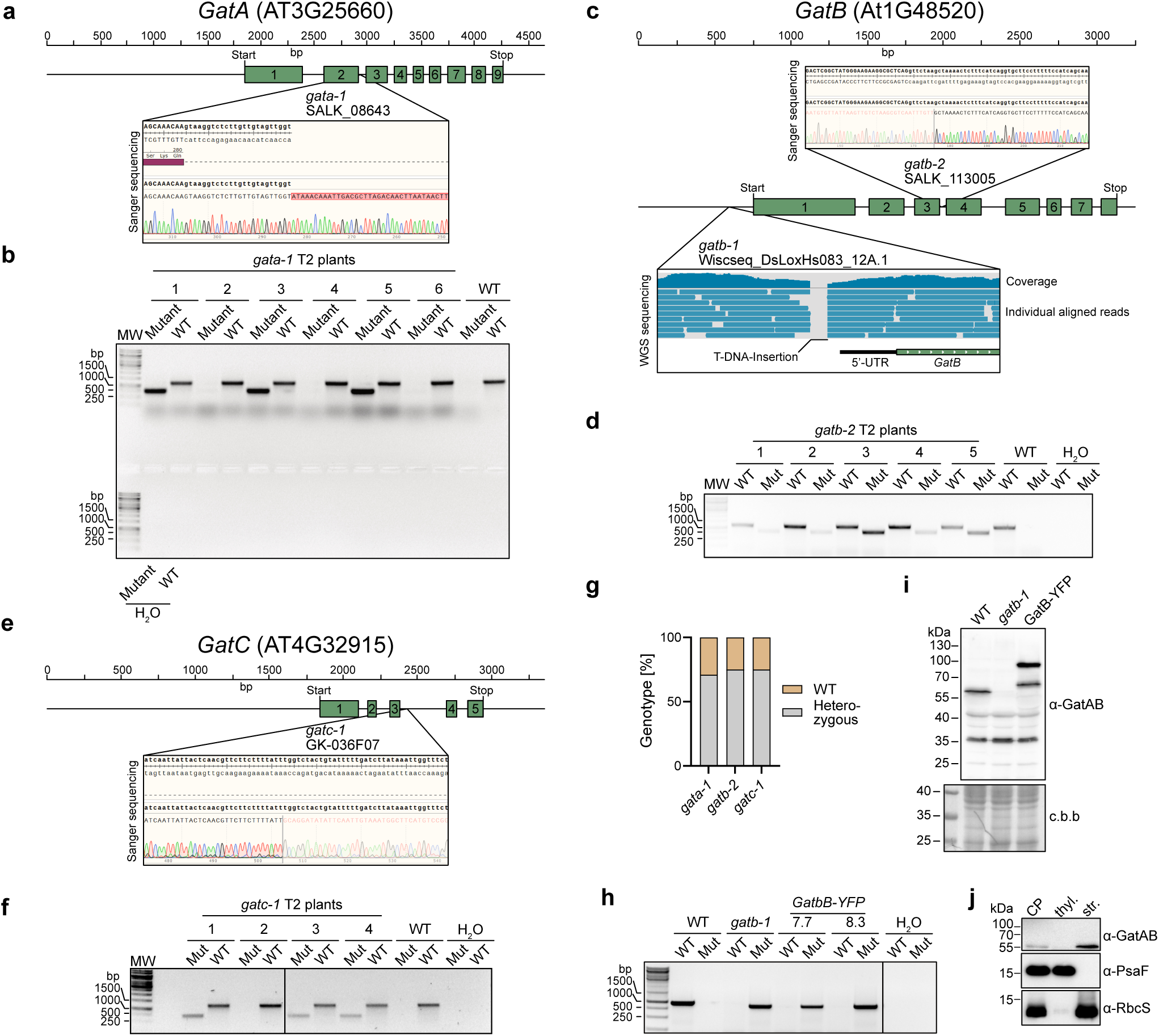
Characterization of *GatA*, *GatB* or *GatC*: T-DNA insertion lines, antibody specificity and sub-organellar localization. (a, c and e) Mapping of the exact T-DNA insertion site of that *gata-1* (a), *gatb-1*, *gatb-2* (c), or *gatc-*(e). Gene structures are illustrated by green introns. The T-DNA insertion sites of *gata-1, gatb-*or *gatc-1* were determined by sanger sequencing of PCR products amplifying the insertion border region of the T-DNA and the adjacent gDNA. The T-DNA insertion site in the *gatb-1* mutant was confirmed by whole genome sequencing of the outcrossed T2 pool of plants. (b, d, f and h) Representative DNA gel electrophoresis images showing the genotyping PCRs for *gata-1* (b), *gatb-2* (d), *gatc-1* (f) and *gatb-1* (including the complementation lines; h). (g) Percentage of genotypes in the T2 generation of *gata-1*, *gatb-2* and *gatc-1* determined by genotyping PCR. n=37-42. (i) Western blot analysis of whole leaf extracts of WT, *gatb-1* and *gatb-1*-GatB-YFP plants to determine the specificity of the α-GatAB antibody. After immunodetection the same membrane was stained with coomassie brilliant blue (c.b.b) which serves as loading control. (j) Immunoblot analysis of whole chloroplasts (CP), thylakoid membranes (thyl.) and stroma (str.) extracts from WT plants probing the localization of the native GatCAB complex using the α- GatAB antibody. α-PsaF and α-Rbcs antibodies serve as marker for thylakoid membranes and stroma, respectively.

**Extended Data Fig. 2:**
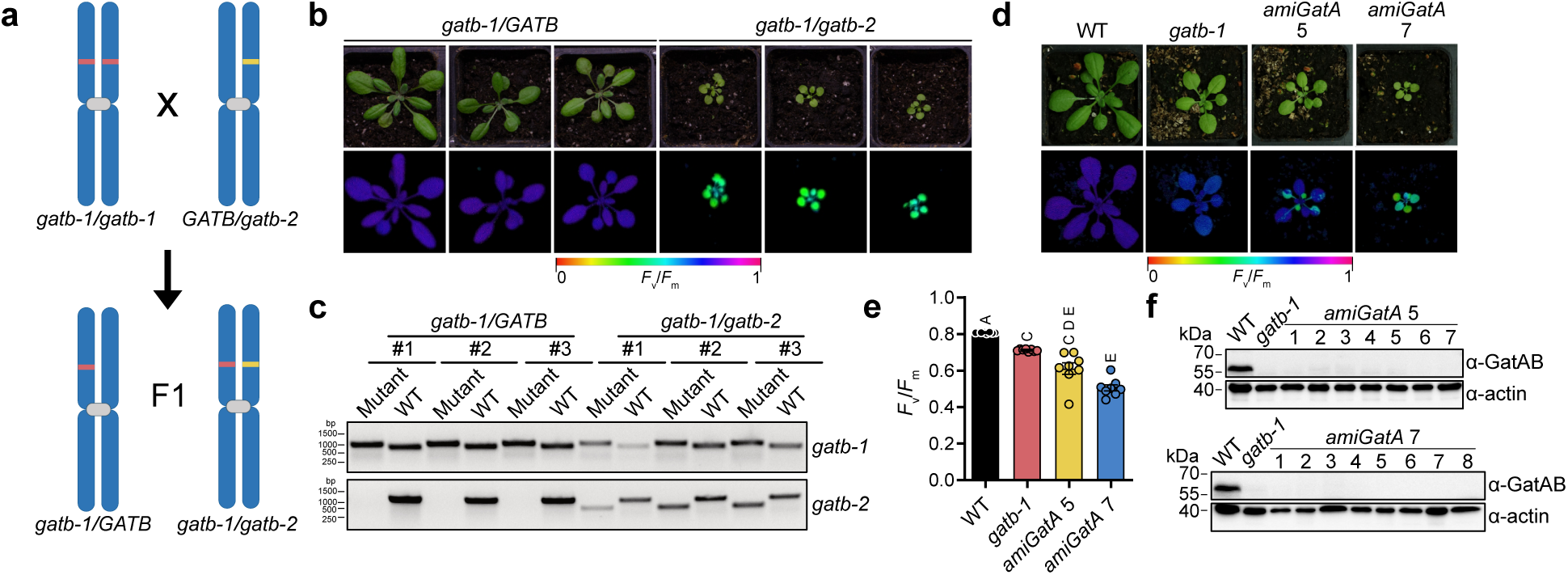
Characterization of *gatb-1gatb-2* and amiRNA GatA (*amiGatA*) mutant plant lines. (a) Schematic representation of the distribution of the *gatb-1* and *gatb-2* mutant alleles on the chromosomes for the crossing of the homozygous *gatb-1* and heterozygous *gatb-2* plants. (b) Photographs and PAM images (*F*_v_/*F*_m_) of representative plants of the first generation after crossing homozygous *gatb-1* and heterozygous *gatb-2* plants. (c) Genotyping PCRs determining the genotype of the plants shown in panel (b). The upper row shows genotyping of the *gatb-1* mutation while the lower one depicts the assessment of the *gatb-2* genotype. (d) Photographs and PAM images (*F*_v_/*F*_m_) of representative plants of two independent lines transformed with an amiRNA targeting the GatA gene (*amiGatA* line 5 and 7) in the second generation after transformation (T2) together with WT and *gatb-1*. (e) Bar graphs of the *F*_v_/*F*_m_ values of WT, *gatb-1*, *amiGatA* line 5 and *amiGatA* line 7. Mean +/- SEM; n=8; Ordinary one-way ANOVA. Letters indicate significantly different groups. (f) Western Blot analysis of the plants used in panel (d) and (e) probing for the presence of GatAB using the α-GatAB antibody. Near equal loading was controlled by using an antibody against actin (α-actin) on the same membranes.

**Extended Data Fig. 3:**
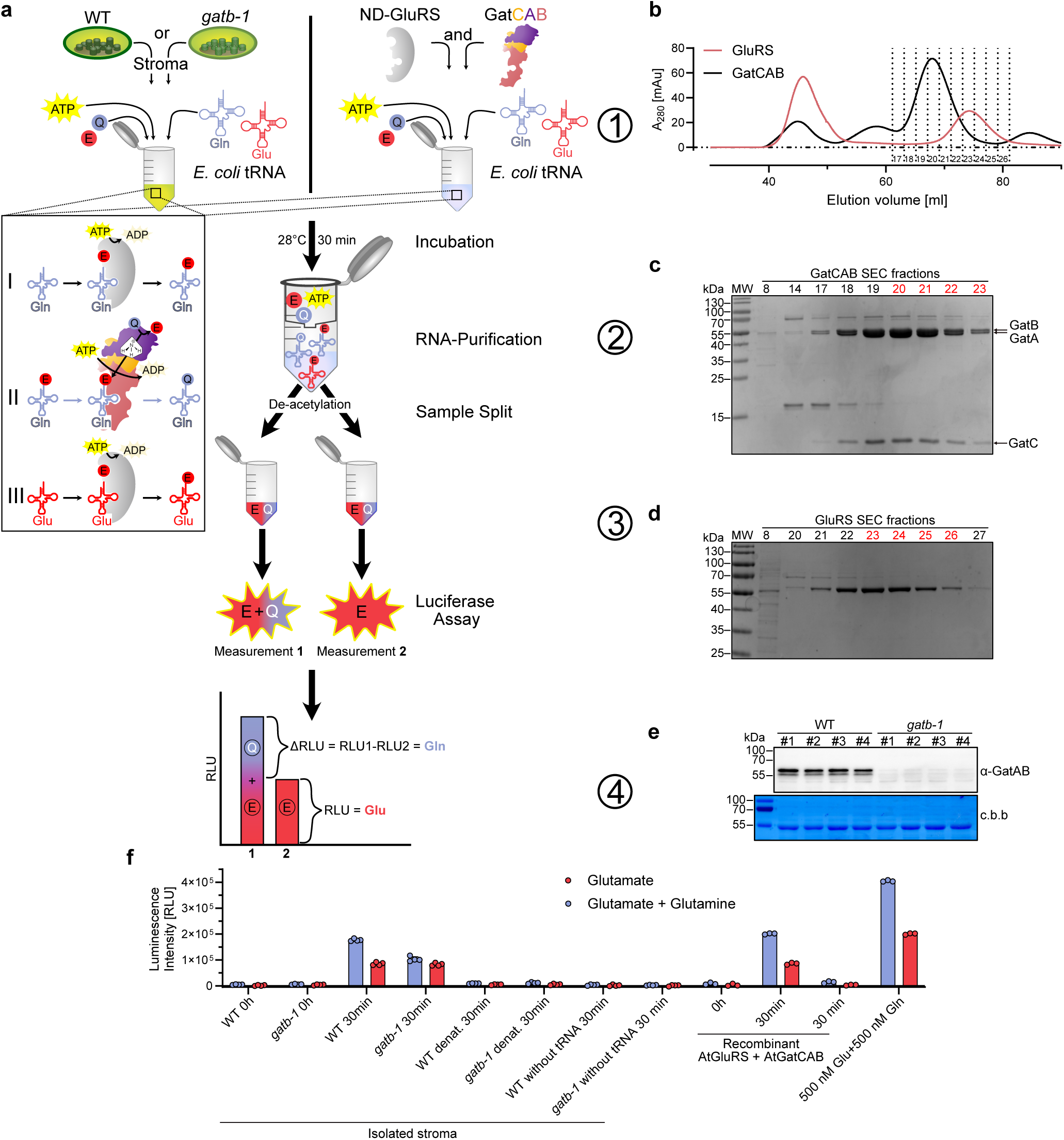
Radioactive label-free amino-acylation and t-RNA-dependent trans- amidation assay with recombinant GluRS + GatCAB and isolated stroma from WT and *gatb-1* plants. (a) Schematic workflow of the label free quantitative assay assessing amino-acylation and transamidation rates in WT and *gatb-1* stroma, compared to recombinant ND-GluRS and GatCAB by a coupled Luciferase Assay. (1) Isolated stroma or recombinant proteins were mixed with ATP, glutamate, glutamine (amine donor) and total E. coli tRNA, followed by incubation. The box illustrated the reactions taking place when stroma was added: (I) Aminoacylation of tRNA^Gln^ with Q by the ND-GluRS; (II) Trans-amidation of Q-tRNA^Gln^ by the GatCAB complex resulting in E- tRNA^Gln^; (III) Aminoacylation of tRNA^Glu^ with Q by the ND-GluRS (2) The reaction was stopped, RNA was purified, and tRNAs were deacetylated to release the covalently bound amino acids. (3) The sample was divided into two separate tubes and a coupled luciferase assay was conducted on each sample to measure the combined glutamine (blue) and glutamate (red) signal in Measurement 1 and the glutamate-only (red) signal in Measurement 2. (4) Glutamine-only (blue) levels were calculated by subtracting the glutamate-only (red) signal from the combined signal. (b) Size exclusion chromatography (SEC) of recombinantly expressed and isolated Arabidopsis proteins ND-GluRS (red) and GatCAB (black) using HiLoad 16/60 Superdex 200 pg column. Dashed lines represent the collected fractions, with the corresponding fraction number indicated at the bottom. (c) SDS-PAGE analysis of GatCAB gel filtration fractions stained with Coomassie blue. Fractions 20-23 (red) were combined and used for further analyses. (d) SDS-PAGE analysis of ND-GluRS gel filtration fractions stained with Coomassie blue. Fractions 23-26 (red) were combined and used for further analyses. (e) Western Blot analysis of normalized WT and *gatb-1* stromal replicates used for the label free enzymatic assay (panel f) detecting GatAB levels using the α-GatAB antibody. After immunodetection the same membrane was stained with coomassie brilliant blue (c.b.b) which serves as loading control. (f) Raw luminescence intensities of the label free quantitative biochemical assay conducted on isolated stroma from WT and *gatb-1* plants, as well as on samples containing recombinant ND- GluRS + GatCAB protein. The data compares glutamate levels (red) and the combined levels of glutamate plus glutamine (blue).

**Extended Data Fig. 4:**
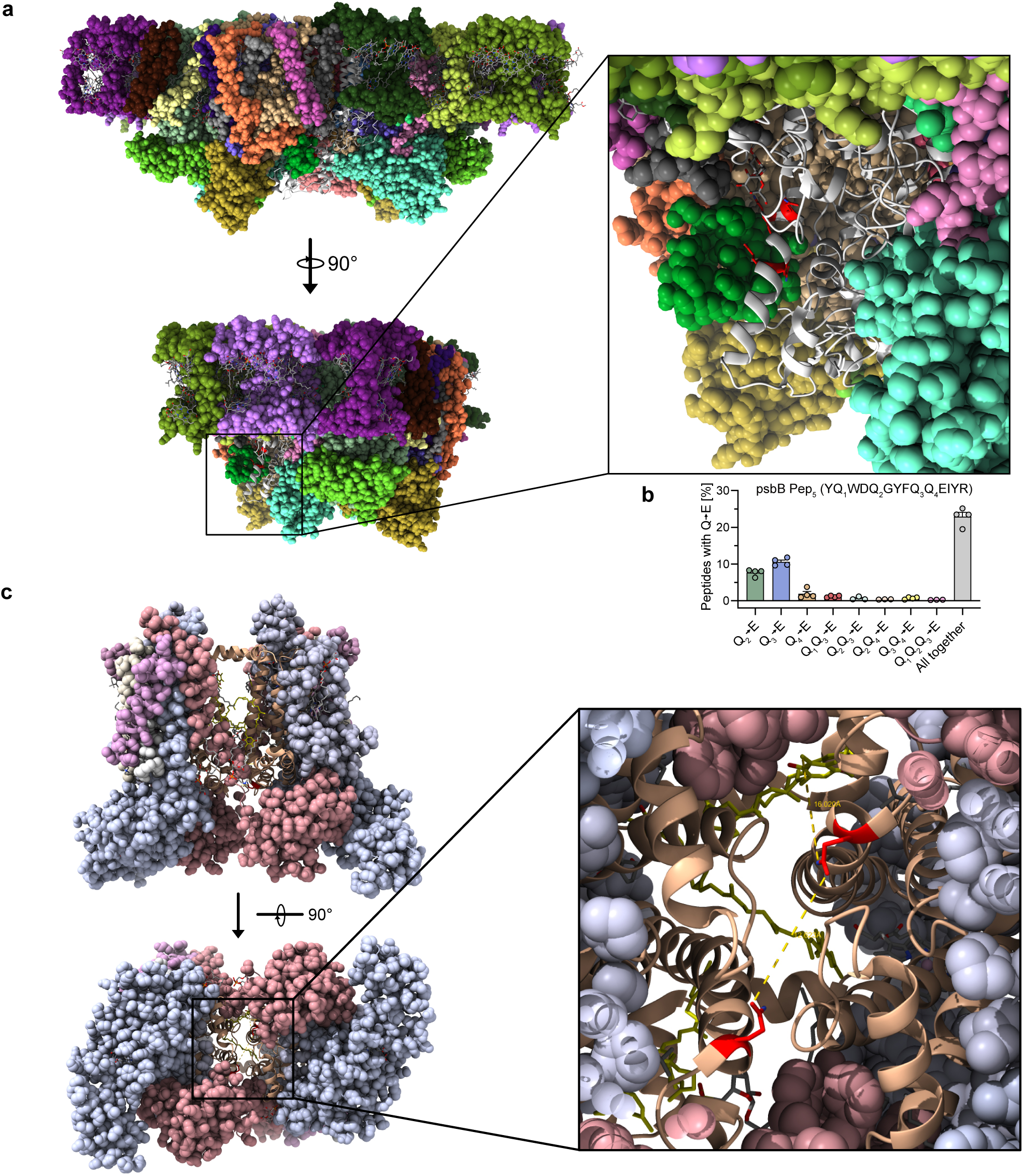
Mapping of peptides of interest on the structure of the photosystem II and cytochrome *b*_6_*f* protein complexes. (a) Mapping of the psbB Pep5 onto the photosystem II complex structure. Sphere (also called Corey-Pauling-Koltun, CPK) representation of the fully assembled spinach photosystem II (PDB-ID: 3jcu). The psbB subunit is shown in ribbons with the Qs of interest in red sticks. The zoom in window shows a close up of the localization of psbB peptide 5. (b) Q→E misincorporation rate of individual peptides (in%) in psbB Pep5 in *gatb-1* measured by shotgun proteomics of isolated chloroplasts. Please note that the individual values are also shown in Fig. 2e. (c) Localization of petB Pep1 in the cytochrome *b*_6_*f* complex. The cytochrome *b*_6_*f* complex (PDB- ID: 6rqf) except petB is shown in the sphere (also called Corey-Pauling-Koltun, CPK) representation. The petB subunit is depicted as gold ribbons. Q177 in Pep1 is shown as red sticks while plastoquinone is illustrated as yellow sticks. Distances between the two opposing Q177 and of Q177 with the adjacent plastoquinone are indicated in the zoomed in box (yellow dashed lines).

**Extended Data Fig. 5:**
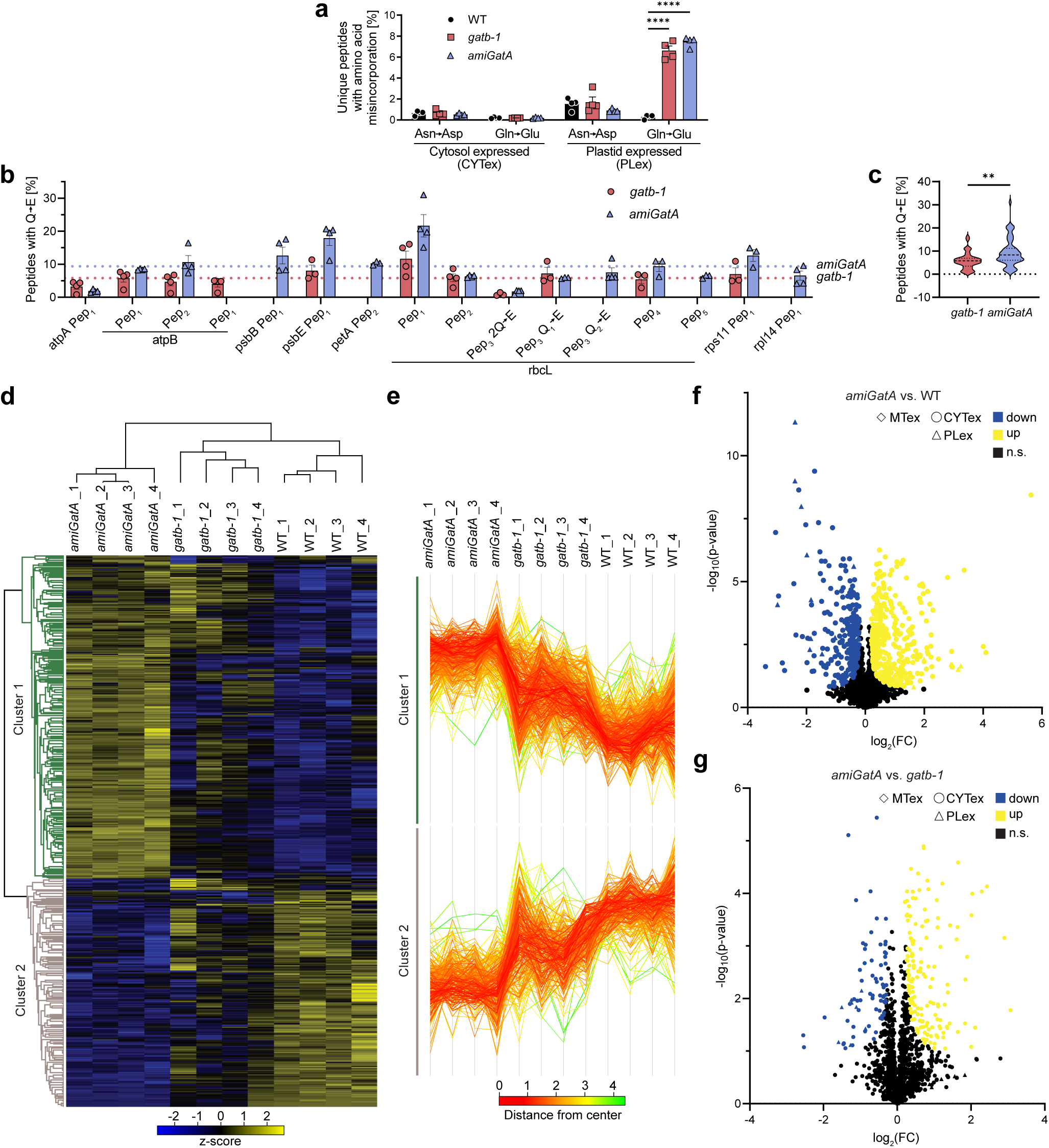
Q→E amino acid misincorporation and changes in the whole leaf proteome in *amiGatA* and *gatb-1* plants. (a) Percentage of unique peptide with either N→D or Q→E substitutions in either cytosolic or plastid expressed proteins in WT (black bars), *gatb-1* (red bars) or the plant expressing the amiRNA against *GatA* (*amiGatA*, blue bars) lines measured by whole leaf shotgun proteomics. The number of unique peptides was divided by the number of unmodified peptides of either CYTex or PLex per biological replicate. Mean +/- SEM; n= 4; Two-way ANOVA with Tukey correction for multiple comparisons. Adjusted p-values ≤ 0.05 are indicated by asterisks. ****: adj. p-value ≤ 0.0001. (b) Q→E misincorporation rate of individual peptides of chloroplast expressed proteins (in%) in *gatb-1* (red bars) and *amiGatA* (blue bars) measured by whole leaf shotgun proteomics. The mean Q→E mis-incorporation rates (see panel c) are shown by dashed lines for *gatb-1* (red) and *amiGatA* (blue). (c) Q→E misincorporation rates (in%) of all peptides PLex in *gatb-1*(red) or *amiGatA* (blue) shown as violin plot (dashed line and dotted lines represent the median and the upper and lower quartiles, respectively). Unpaired t-test with Welch’s correction; **: p-value ≤ 0.01 (d) Heat map showing the row z-score of significantly changed proteins (ANOVA with permutation-based FDR of 5%) in WT, *gatb-1* and *amiGatA* leafs measured by shotgun proteomics. Both rows and columns were hierarchically clustered (Euclidian with average linkage). (e) Cluster profile plots of the two main cluster of DEPs (d). Each line represents the z-score of one protein and the color represents the distance from the cluster center. (f-g) Volcano plots showing differentially expressed proteins (DEPs) in *amiGatA* when compared to WT (f) or *gatb-1* (g) (FDR = 5% and S0 = 0.1). The log2 of the fold change (FC) is plotted against the negative log10 of the adjusted p-value. Significantly up-regulated genes are colored yellow and down-regulated genes a shown in blue while non-significantly regulated proteins (n.s.) are depicted in black. Cytosolic expressed proteins are indicated by circles, plastid expressed proteins by triangles and mitochondrially expressed proteins by diamonds.

**Extended Data Fig. 6:**
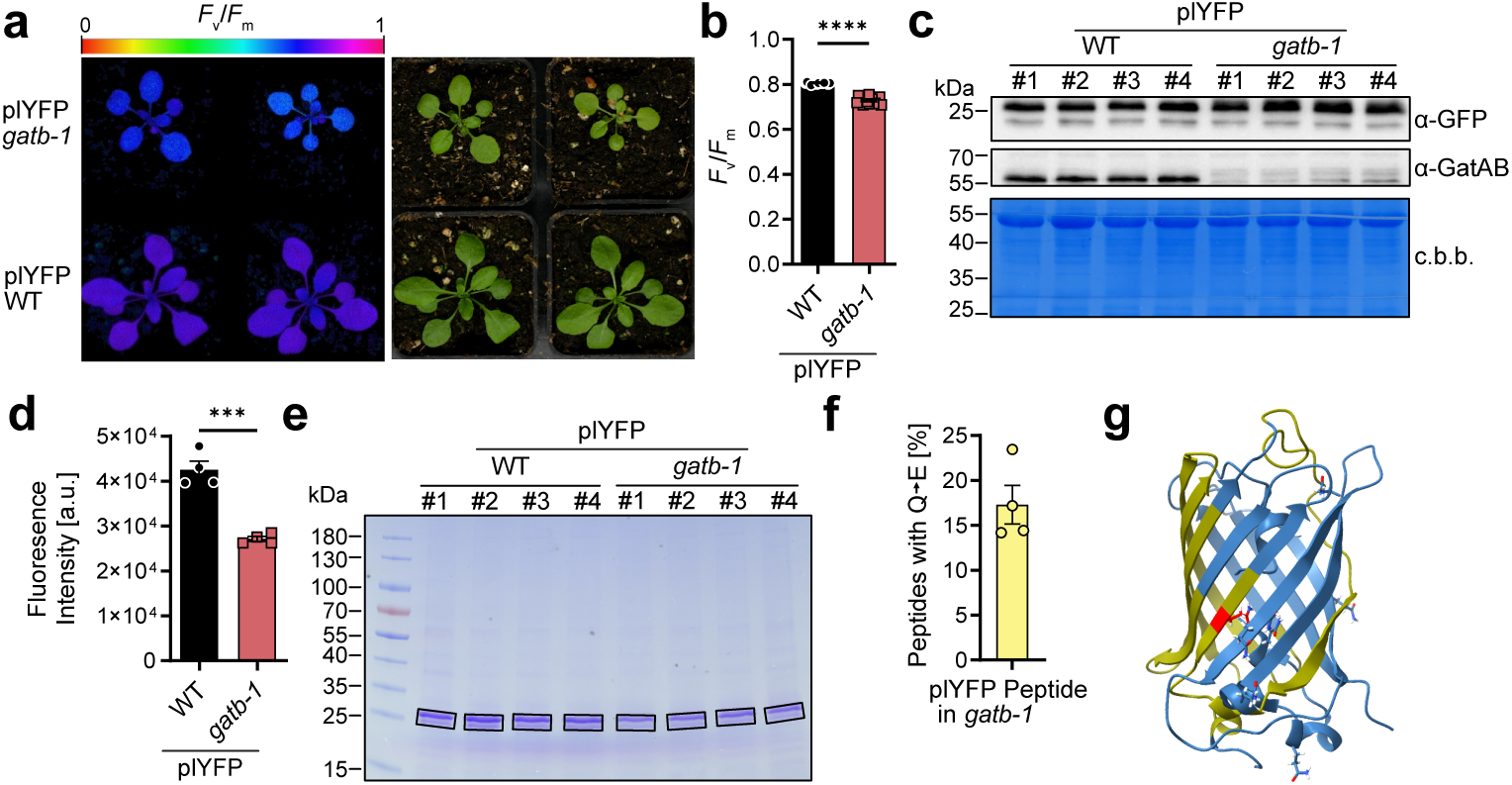
Plastid expressed YFP in the *gatb-1* genetic background results in reduced YFP fluorescence due to Q→E misincorporation in YFP. (a) Photographs and PAM images (*F*_v_/*F*_m_) of representative plants of homozygous T3 plants with plastid expressed YFP (plYFP) in the WT (plYFP WT, bottom) and or *gatb-1* (plYFP *gatb-1;* top) background. (b) Bar graphs of the *F*_v_/*F*_m_ values of plYFP WT (black bars) and plYFP *gatb-1* (red bars). Mean +/- SEM; n= 18; Unpaired t-test with Welch’s correction. ****: p-value ≤ 0.0001. (c) Immunoblot showing YFP amounts (α-GFP antibody), presence of GatAB (α- GatAB) and equal loading (coomassie-stained membrane, c.b.b.) of the extracts used to determine the fluorescence intensity in (c). (d) YFP fluorescence intensity of total leaf protein extracts of plYFP WT (black bars) and plYFP *gatb-1* (red bars). Mean +/- SEM; n= 4; Unpaired t-test. *** = p-value ≤ 0.001 (e) Gel image showing immuno-precipitated YFP from plYFP WT and plYFP *gatb-1* plants used for the proteomics analysis. The black boxes indicated bands cut out and subjected to the mass spectroscopic analysis. (f) Q→E misincorporation rate of the Q containing peptides of chloroplast expressed YFP immuno- precipitated from plYFP *gatb-1* plants (in%) measured by mass spectrometry of the gel slices shown in (F). No mutant peptides were found YFP precipitated from plYFP WT plants. Mean +/- SEM; n= 4 (g) YFP 3D structure shown as ribbons indicating the Q which is substituted to E in *gatb-1*. Parts of the protein covered by the MS analysis are indicated in yellow and the Q which is changed to E in the plYFP peptide is shown as stick in red.

**Extended Data Fig. 7:**
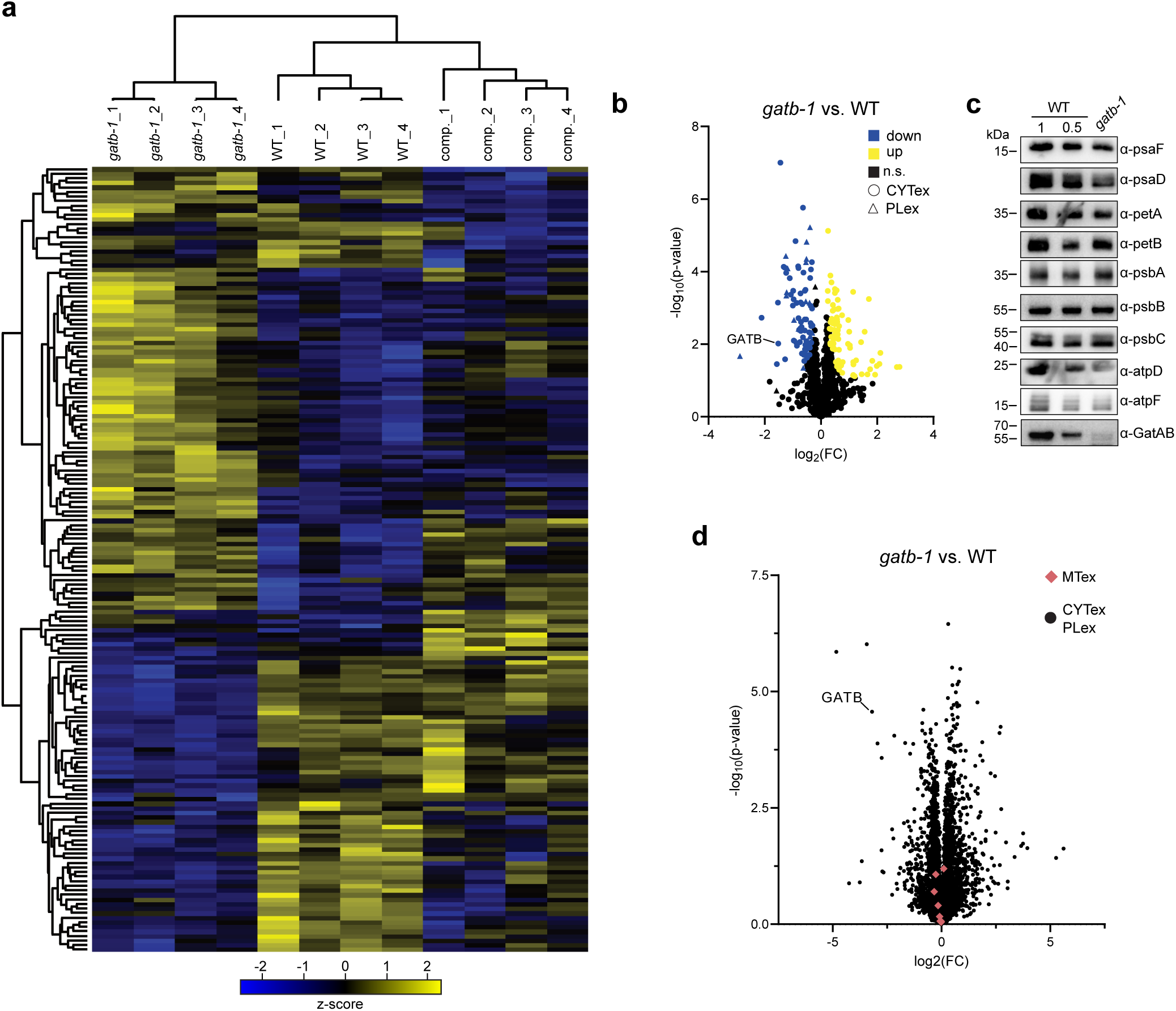
Label-free quantification of chloroplast proteins for WT, *gatb-1* and comp., LFQ for WT and *gatb-1* mitochondrial proteins and western blot confirmation of selected proteins. (a) Heat map showing the row z-score of significantly changed proteins (ANOVA with permutation-based FDR of 5%) in WT, *gatb-1* and complementation chloroplasts. Both rows and columns were hierarchically clustered (Euclidian with average linkage). (b) Volcano plot showing differentially expressed proteins (DEPs) in *gatb-1* when compared to WT (FDR = 5% and S0 = 0.1). The log2 of the fold change (FC) is plotted against the negative log10 of the p-value. Significantly up-regulated genes are colored yellow and down-regulated genes a shown in blue while non-significantly regulated proteins (n.s.) are depicted in black. Cytosolic expressed proteins are indicated by circles and chloroplast expressed proteins by triangles. (c) Western blot of selected proteins using the same chloroplasts than used for the shotgun mass spec analysis. Loaded protein amounts were normalized. Of WT chloroplast extracts the same amount (1) and half of the amount (0.5) of *gatb-1* proteins were loaded for easier comparison. (d) Volcano plot showing differentially expressed proteins (DEPs) in *gatb-1* when compared to WT (FDR = 5% and S0 = 0.1). The log2 of the fold change (FC) is plotted against the negative log10 of the p-value. Mitochondrially expressed proteins are shown as red diamonds while proteins expressed in the cytosol and plastids are depicted as black circles.

**Extended Data Fig. 8:**
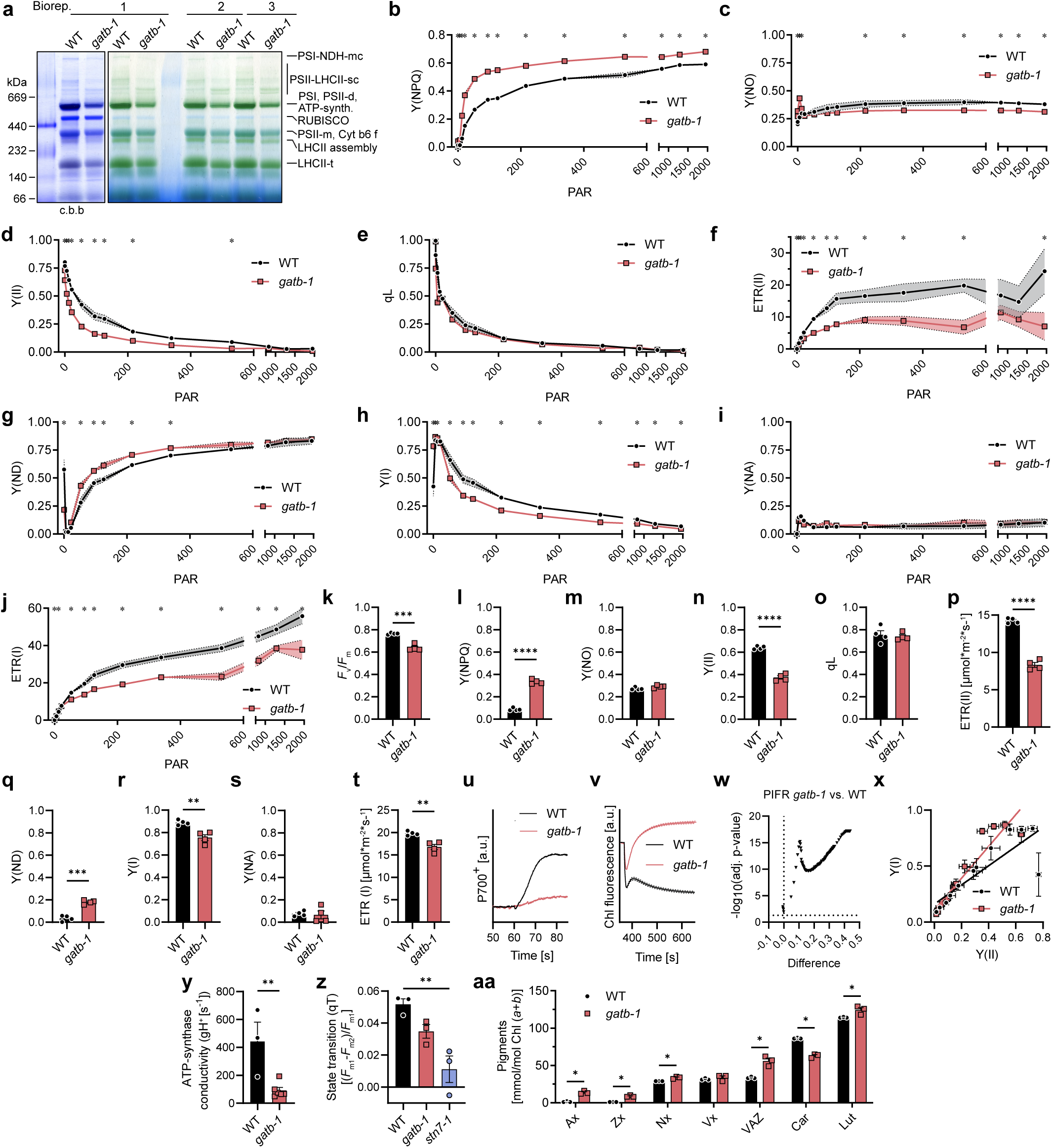
The *gatb-1* mutation results in slightly changed assemblies of photosynthetic complexes and photosynthetic parameters. (a) BN-PAGE analysis of all bio replicates of WT and *gatb-1* chloroplasts to assess the assembly of photosynthetic complexes. The amount of loaded chlorophyll was normalized to the fresh weight of the plants (see material and methods). For biological replicate 1, the same area of the gel is depicted before (right) and after (left) Coomassie staining (c.b.). Please note that biological replicate 1 is also shown in Fig. 3a. mc: mega complex; sc: supercomplex; m: monomer; t: trimer. (b-j) Photosynthetic parameters of WT (black circles) and *gatb-1* (red squares) plants measured at different light intensities. Mean +/- SEM; n= 5; Multiple unpaired t-tests with correction for multiple testing according to Benjamini, Krieger, and Yekutieli using a false discovery rate (FDR) of 5%. A single asterisk above a data point indicates a significant difference of with the indicated FDR. (k-t) Photosynthetic parameters of WT (black bars) and *gatb-1* (red bars) plants measured analysis at 53 PAR. Mean +/- SEM; n= 4. Unpaired t-test; ** = p-value ≤ 0.01, *** = p-value ≤ 0.001, **** = p-value ≤ 0.0001. (u) Post illumination fluorescence signal (PIFR) of WT (black lines) and *gatb-1* (red lines) plants. Mean +/- SEM; n= 9. Multiple unpaired t-tests with correction for multiple testing according to Benjamini, Krieger, and Yekutieli using a false discovery rate (FDR) of 5%. Every *gatb-1* datapoint was significantly different than WT (see volcano plot in Extended Data Fig. 8l) (v) Volcano Plot of the multiple t-tests of the post illumination fluorescence signal shown in Fig. 3I. Each point represents the result of one t-test. (w) Photosystem II P700 oxidation measurements of WT (black lines) and *gatb-1* (red lines) plants. (x) Y(II) values plotted against the Y(I) values for WT (black circles) and *gatb-1* (red squares) for different light intensities as measured in panel (d) and (h). Linear regression lines are shown (WT: black, *gatb-1*: red) and are significantly different test by an equivalent of an analysis of covariance (ANCOVA). Mean +/- SEM; n= 5. (y) ATP synthases activity WT (black bars) and *gatb-1* (red bars) plants. Mean +/- SEM; n= 3-6. Unpaired t-test; ** = p-value ≤ 0.01. (z) State-Transition measurements of WT (black bars), *gatb-1* (red bars) and the state transition mutant *stn7-1* (blue bars). Mean +/- SEM; n= 3; Ordinary one-way ANOVA. ** = adj. p-value ≤ 0.01. (aa)Pigment content of WT (black bars) and *gatb-1* (red bars) normalized to chlorophyll a and b. Ax = antheraxanthin, Zx = zeaxanthin, Nx = neoxanthin, Vx = violaxanthin, VAZ = sum of Vx, Ax and Zx and Car = β-carotene. Multiple unpaired t-tests with correction for multiple testing according to Benjamini, Krieger, and Yekutieli using a false discovery rate (FDR) of 5%. A single asterisk indicates a significant difference with the indicated FDR.

**Extended Data Fig. 9:**
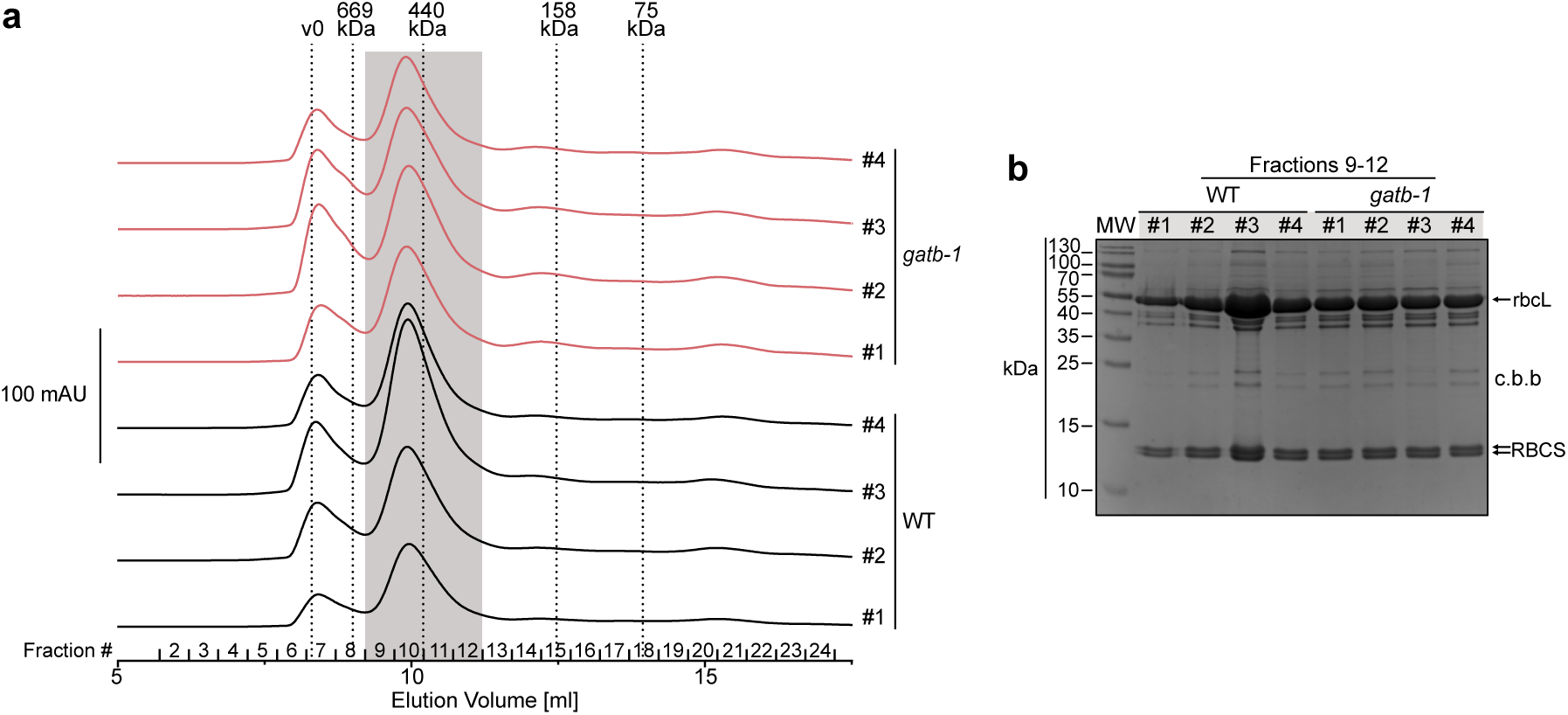
Size exclusion chromatography and SDS-PAGE of the isolation of fully assembled L_8_S_8_ RuBisCO. (a) Size exclusion chromatographs (SEC) of all isolations of L_8_S_8_ RuBisCO from WT or *gatb-1* stroma. Fractions 9-12 (gray background) were combined. (b) Coomassie stained SDS-PAGE of the combined fractions of the SEC runs shown in (a).

**Extended Data Fig. 10:**
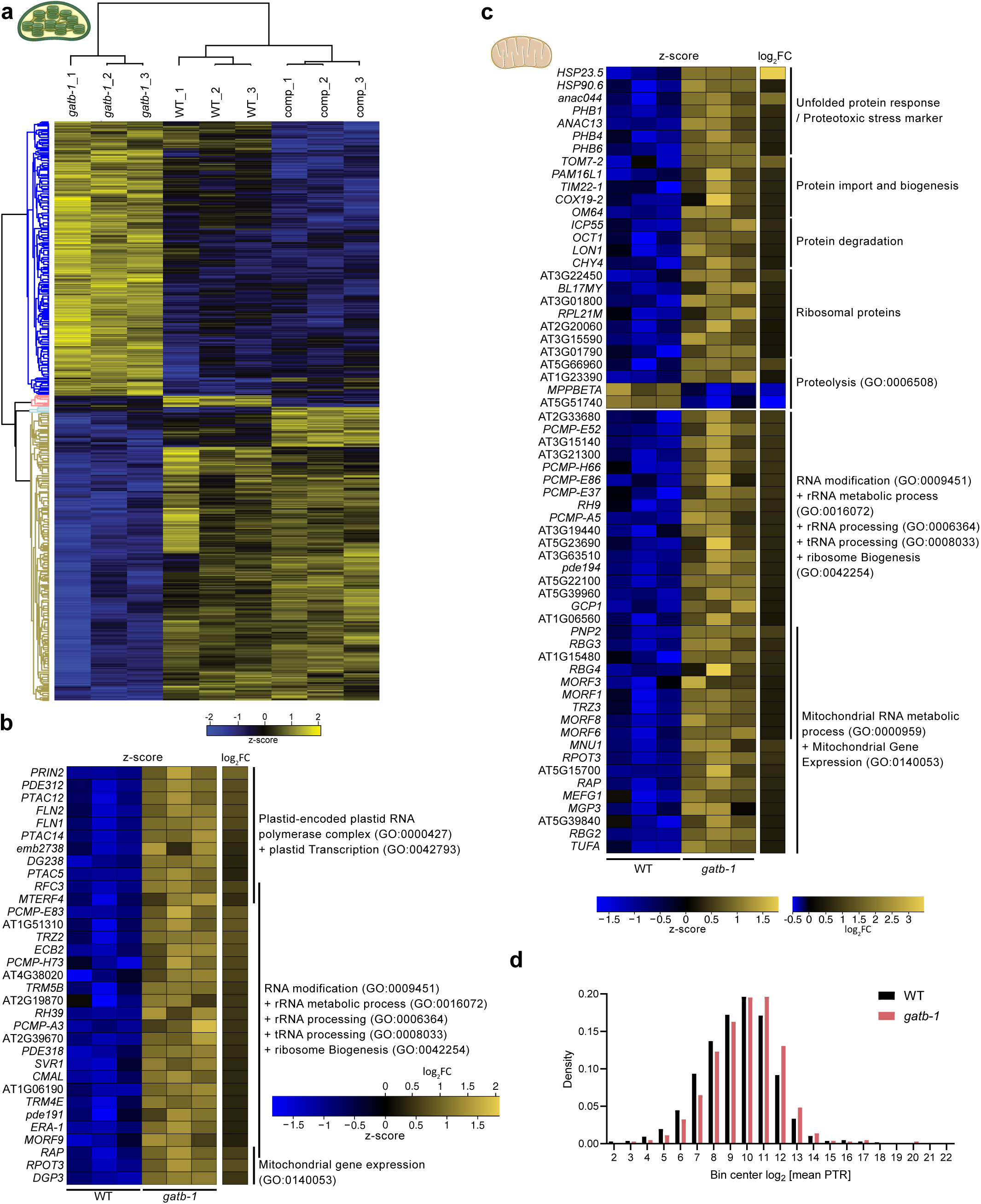
Transcriptomics of WT, *gatb-1* and comp. lines reveals compensatory mechanisms. (a) Heat map showing the row z-scores of significantly changed genes (ANOVA with permutation-based FDR of 5%) in WT, *gatb-1* and comp. plants measured by mRNA sequencing. Both rows and columns were hierarchically clustered (Euclidian with average linkage). (b) Heatmap depicting the row z-scores and the log2FCs of a selection of significantly changed genes (adjusted p-value <0.05) coding for chloroplast localized proteins (according to SUBA) which are included in interesting and significantly enriched GO-terms. Some genes have overlapping assignments. (c) Heatmap depicting the row z-scores and the log2FCs of a selection of significantly changed genes (adjusted p-value <0.05) coding for mitochondria localized proteins which are either hand-curated (up) or included in interesting and significantly enriched GO-terms (bottom). Some genes have overlapping assignments to GO-terms. (d) Distribution of protein to transcript ratios (PTRs) of WT and *gatb-1* proteins/genes.

**Extended Data Fig. 11:**
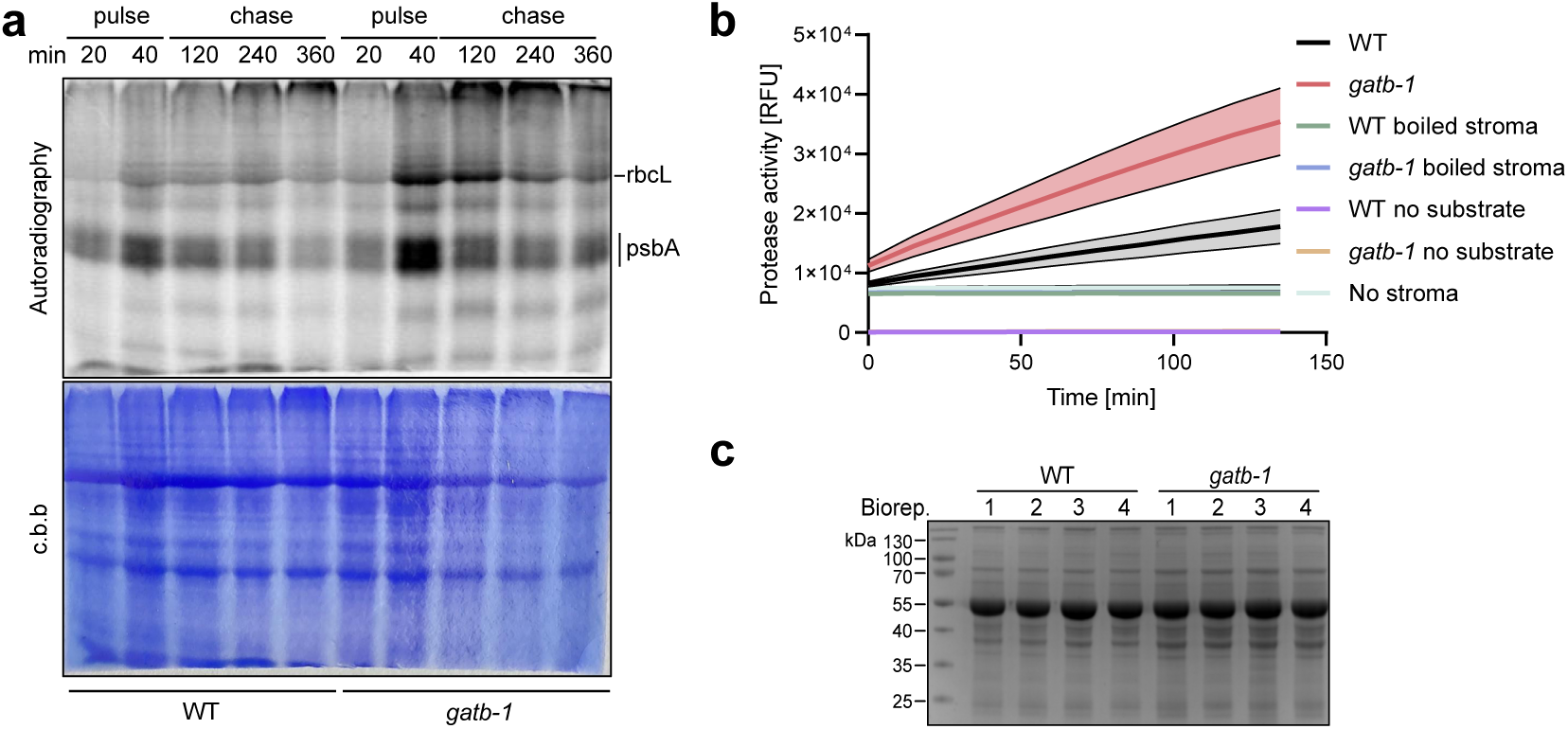
Representative gel of *in vivo* radiolabeling assay and complete data of protease assay. (a) Representative gel of *in vivo* radiolabeling assay of WT (left) and *gatb-1* (right) assays. Radioactive amino acids were added at time point 0, removed after 40 minutes of pulse and then incubated an additional 320 minutes of chase. Samples were taken and subjected to SDS-PAGE at the indicated time points. The prominent bands of the RuBisCO large subunit (rbcL) and of the A subunit of photosystem II (psbA) in the autoradiography are indicated. The lower panel show the same gel stained with coomassie brilliant blue (c.b.b). (b) Raw data and controls for the assay assessing the protease activity of WT (black, green and violet lines) and *gatb-1* (red, blue and ochre lines). Mean +/- SEM; n= 4. Please note that the protease activity rates shown in Fig. 5g were calculated from the data in this Fig.. (c) SDS-PAGE of the stromal extracts used in the protease assay shown in panel (b) illustrating equal loading.

